# Orbitofrontal Cortex Influences Dopamine Dynamics Associated with Alloparental Behavioral Acquisition in Female Mice

**DOI:** 10.1101/2023.02.03.527077

**Authors:** Gen-ichi Tasaka, Mitsue Hagihara, Satsuki Irie, Haruna Kobayashi, Kengo Inada, Kenta Kobayashi, Shigeki Kato, Kazuto Kobayashi, Kazunari Miyamichi

## Abstract

Maternal behaviors, which are crucial for the survival of mammalian infants, require the coordinated operation of multiple brain regions to process infant cues, make decisions, and execute motor plans. Although these processes likely demand higher cognitive functions, the prefrontal areas that regulate limbic parental programs remains poorly understood. Here, we show that the orbitofrontal cortex (OFC) excitatory projection neurons promote alloparental caregiving behaviors in female mice. By chronic microendoscopy, we observed robust yet adaptable representations of pup-directed anticipatory and motor-related activities within the OFC. Some of these plastic responses were significantly overlapped with those related to nonsocial reward signals. The inactivation of OFC output reduced the phasic activities of midbrain dopamine (DA) neurons specifically tied to pup retrieval and impaired the modulation of DA release to the ventral striatum during the acquisition of alloparental behaviors. These findings suggest that the OFC transiently boosts DA activity during the acquisition phase, thereby facilitating the manifestation of alloparental behaviors.

## Introduction

Maternal behaviors, including feeding and protection of the young, are crucial for the physical and mental health of mammalian infants (*1*). To achieve these behaviors, many brain areas must work together to process infant cues, make decisions, and generate motor planning and execution, which likely requires higher cognitive function of the brain. Although classical models of neural circuits for parental behaviors often assume the prefrontal cortex (PFC) as an integrator of infant-related sensory signals and a controller of decision-making and motivation (*1, 2*), these ideas are mostly based on anatomical studies and have not been functionally tested. On one hand, maternal behaviors are triggered by a pup’s sensory signals, and a large body of research has examined maternal plasticity in sensory coding in the olfactory and auditory regions (*3-8*). On the other hand, classical lesion and pharmacological manipulation studies in rats have emphasized the importance of the medial preoptic area and its downstream mesolimbic dopamine (DA) system in maternal caregiving motivation (*9*). Recent molecular genetic studies in mice have described more precise circuit architectures of subcortical parental behavioral centers at the resolution of specific cell types (*10-14*). In contrast to these advances in understanding cortical sensory systems and subcortical parental behavioral centers, the functional organization of the PFC that potentially links sensory signals to maternal behavioral execution remains mostly elusive (*15*). As such, little is known about how subcortical regions are regulated by the cortical top-down inputs in the context of maternal behaviors.

There is accumulating evidence that the PFC is involved in many aspects of social behaviors, such as social dominance (*16, 17*), sexual behaviors (*18*), and social observational fear learning (*19, 20*). Although most previous studies have focused on the medial PFC (mPFC), a recent study reported that the orbitofrontal cortex (OFC) of mother mice is activated by the ultrasonic vocalizations of pups (*4*). Imaging studies in humans have also shown that visual and auditory signals related to children evoke activity in the OFC of mothers (*21, 22*). The OFC is generally thought to facilitate the transition between sensation and action by associating diverse sensory signals with valence in behavioral contexts (*23-26*). Furthermore, recent studies have revealed that individual OFC neurons encode diverse representations of behaviorally relevant features, including sensory signals, spatial locations, and cues that predict reward outcomes, as well as rewards *per se* (*27-29*). However, the neural representations or functions of the OFC in guiding naturally occurring behaviors, such as maternal caregiving behaviors, remain unexplored. This knowledge gap motivated us to explore the functional representations of pup-directed caregiving behaviors in the OFC of female mice.

Mothers are innately motivated to care for their young with strong rewards (*30*). The ventral tegmental area (VTA) in the midbrain, where most DA neurons are located, is one of the most well-characterized regions for reinforcement learning (*31, 32*). The neural activity of DA neurons in the VTA (VTA^DA^ neurons) ramps up during pup retrieval (*11*), a hallmark of maternal caregiving behaviors, and gradually decreases as female mice experience more pup retrieval (*33*). This dynamics of VTA^DA^ neuron activity is thought to reflect reward prediction error (RPE) to guide reinforcement learning (*33*), although the precise neural mechanisms underlying such plasticity remain largely unknown. Pharmacological or optogenetic inactivation of VTA^DA^ neurons disrupts pup retrieval (*33, 34*), underscoring their crucial role in maternal motivation toward infants. Notably, the OFC projects to the VTA, potentially contributing to reward-based learning (*35-39*). Therefore, we hypothesize that the OFC may modulate VTA^DA^ neurons by conveying pup-related information, which may underlie the motivation for maternal behaviors.

Virgin female mice can exhibit maternal caregiving behaviors, such as pup retrieval, following instruction, observation of experienced caretakers (*40, 41*), or repeated exposure to infants (*42*). We chose to focus on the emergence of maternal retrieval behaviors in virgin female mice (alloparental pup retrieval) for two reasons. First, unlike the rapid onset of maternal behaviors in biological mothers, alloparental retrieval develops gradually and slowly in virgin females (*33, 41*), allowing us to characterize the varying neural correlates of increasing behavioral performance and the dynamics of neural activities during these processes. Second, once maternal behaviors are established, they are robust (*43*). The alloparental paradigm may be particularly suited for characterizing the potentially nuanced modulatory roles of the higher cognitive areas in maternal behaviors. Therefore, we employed a combination of viral-genetic manipulation and chronic microendoscopic calcium (Ca^2+^) imaging to examine the functions of the OFC in the acquisition of alloparental pup retrieval behaviors in virgin female mice.

## Results

### OFC layer 5 neurons are required for the efficient acquisition of alloparental pup retrieval

The OFC is characterized by a thick layer 5 that projects to various subcortical regions (*39*). We utilized the *Rbp4-Cre* line (*44*), a Cre driver specific to excitatory layer 5 neurons, to selectively target excitatory layer 5 pyramidal cells in the OFC (referred to as OFC^Rbp4^ neurons). To examine the role of these neurons in alloparental behaviors, we applied active caspase-mediated genetic cell ablation (*45*) To evaluate ablation efficiency, we injected an AAV encoding taCasp3 into the left hemisphere and phosphate-buffered saline (PBS) into the right hemisphere of *Rbp4-Cre* mice crossed with the *Ai162* reporter line as a marker, which expresses GCaMP6s in a Cre-dependent manner (*46*) (Fig. 1A). This resulted in an approximately 30% reduction in GCaMP6s-expressing (+) cells on the AAV-injected side relative to the PBS-injected control side (Fig. 1B, C). We then bilaterally injected taCasp3 AAV into the OFC and examined the effects of this moderate ablation of OFC^Rbp4^ neurons in the alloparental training paradigm (*41*), in which virgin females were cohoused with a mother and her pups. The retrieval success rates, defined as the fraction of females retrieving all six pups within 4 min, were measured before cohousing and at 1 and 2 days afterward (Fig. 1D) (*3, 41*). The latency to investigate the pups did not significantly differ between groups, suggesting similar levels of attention toward pups (Fig. S1A). Prior to cohousing, a few virgin females showed spontaneous pup retrieval. After two days of cohousing, all females in the control group successfully retrieved all six pups, with a reduced latency (Fig. 1E, F, S1B). This relatively rapid behavioral acquisition, compared to previous studies (*41*), may reflect the genetic background of our study, which utilized F1 hybrids of the B57BL/6J and FVB strains. By contrast, females with ablated OFC^Rbp4^ neurons showed reduced spontaneous retrieval, and five out of twelve failed to achieve successful retrieval, demonstrating a significant impairment in the acquisition of alloparental pup retrieval (Fig. 1E–G).

**Fig. 1.**
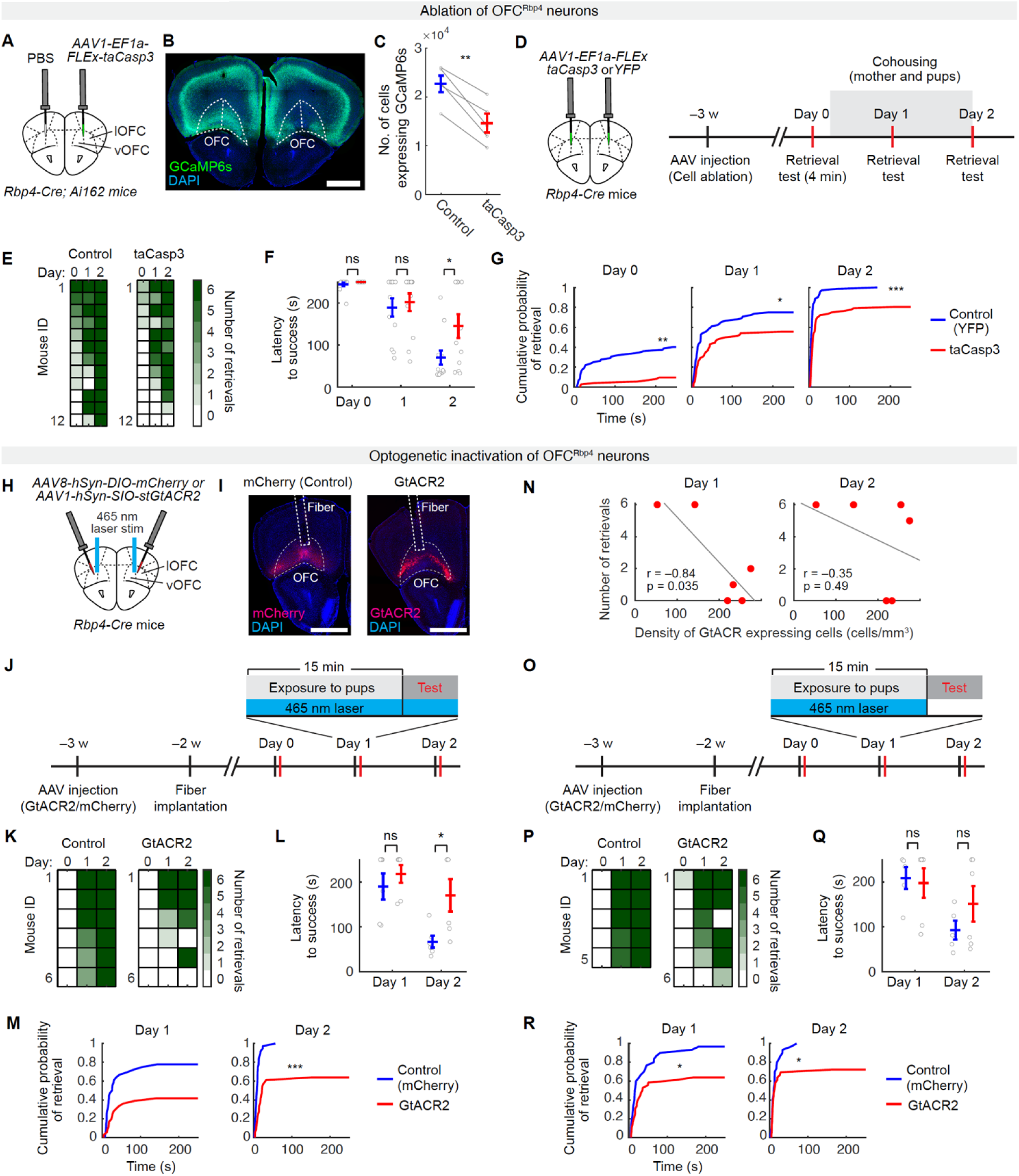
Necessity of OFC^Rbp4^ neurons for effective acquisition of pup retrieval. (A) Schematic of the experimental design. AAV1 *EF1a-FLEx-taCasp3-TEVp* was unilaterally injected into the ventral and lateral OFC (vOFC and lOFC) of *Rbp4-Cre; Ai162* mice. (B) Representative coronal section showing selective ablation in the taCasp3-injected hemisphere. GCaMP6s signals were amplified by anti-GFP staining. Scale bar, 1 mm. (C) Quantification of GCaMP6+ cells in the OFC. **, p < 0.01 by paired t-test (N = 5 mice). (D) Schematic of the experimental design. AAV1 *EF1a-FLEx-taCasp3-TEVp* or AAV1 *EF1a-DIO-YFP* was injected bilaterally in virgin *Rbp4-Cre* mice >3 weeks before testing. (E) Individual pup retrieval performance. (F) Latency to retrieve all six pups in a 4-min trial. *p < 0.05 by post-hoc unpaired t-test with Benjamini-Hochberg correction after a significant two-way ANOVA with repeated measures. ns, not significant (N = 12 per group). (G) Cumulative retrieval probability. **, p < 0.01 by Kolmogorov–Smirnov test. (H) Schematic of the experimental design. AAV1 *hSyn-DIO-stGtACR2-fusionRed* or AAV8 *hSyn-DIO-mCherry* was bilaterally injected in virgin *Rbp4-Cre* mice. (I) Representative sections showing the optic fiber tract and mCherry/stGtACR2 expression. Scale bar, 1 mm. (J, O) Schematics of the experimental design. 15-min pup exposure followed by testing with laser during both sessions (J) or cohousing only (O). (K, P) Individual retrieval performance. (L, Q) Retrieval latency. *p < 0.05 by post-hoc unpaired t-test with Benjamini-Hochberg correction after a significant two-way ANOVA with repeated measures. ns, not significant. N = 6 per group in (L); N = 5 (Control) and N = 6 (GtACR2) in (Q). (M, R) Cumulative retrieval probability. *, p < 0.05, and ***, p < 0.001 by Kolmogorov–Smirnov test. (N) Correlation between OFC GtACR2 expression and retrieval performance. Error bars indicate the standard error of the mean (SEM). See Fig. S1 for more data.

To investigate whether OFC^Rbp4^ neuron activity during pup exposure contributes to the induction of pup retrieval, we utilized an inhibitory optogenetic approach coupled with repetitive pup exposure, a protocol known to induce alloparental pup retrieval in virgin female mice (*42*). We administered AAVs encoding the inhibitory opsin GtACR2, with mCherry serving as a control, into the OFC of *Rbp4-Cre* mice (Fig. 1H, I). Optic fibers were then bilaterally implanted above the OFC for optogenetic manipulation (Fig. S1C). Virgin females were exposed to three pups in a small home cage for 15 min per day, followed by a 4-min pup retrieval assay. During both the exposure and retrieval sessions, OFC^Rbp4^ neurons were silenced (Fig. 1J). Prior to pup exposure, no virgin females exhibited spontaneous retrieval behavior, regardless of group (Fig. 1K). While pup investigation latency remained unchanged (Fig. S1D), silencing OFC^Rbp4^ neurons led to fewer successful retrievals and a significantly prolonged retrieval latency (Fig. 1K–M). A negative correlation was observed between the number of GtACR2+ cells and retrieval performance, with a significant correlation on day 1 (Fig. 1N).

To further examine whether OFC activity is required during pup exposure, we implemented a condition in which OFC^Rbp4^ neurons were silenced exclusively during the pup exposure period (Fig. 1O). This manipulation produced a similar but less pronounced reduction in retrieval performance (Fig. 1P, R). Thus, the acute activity of OFC^Rbp4^ neurons during pup exposure is required for facilitating the expression of alloparental pup retrieval behavior through repetitive pup experience.

### Temporal dynamics of pup retrieval-related responses in OFC^Rbp4^ neurons

OFC^Rbp4^ neurons contribute to alloparental pup retrieval, and thus, we next investigated a neural representation of such behaviors by these neurons. We utilized microendoscopic Ca^2+^ imaging (*47*) in *Rbp4-Cre*; *Ai162* female mice, which were surgically implanted with a gradient refractive index (GRIN) lens (500-μm diameter) above layer 5 of the OFC (Fig. 2). The expression of GCaMP6s and the placement of the lens were histologically confirmed (Fig. S2A).

**Fig. 2.**
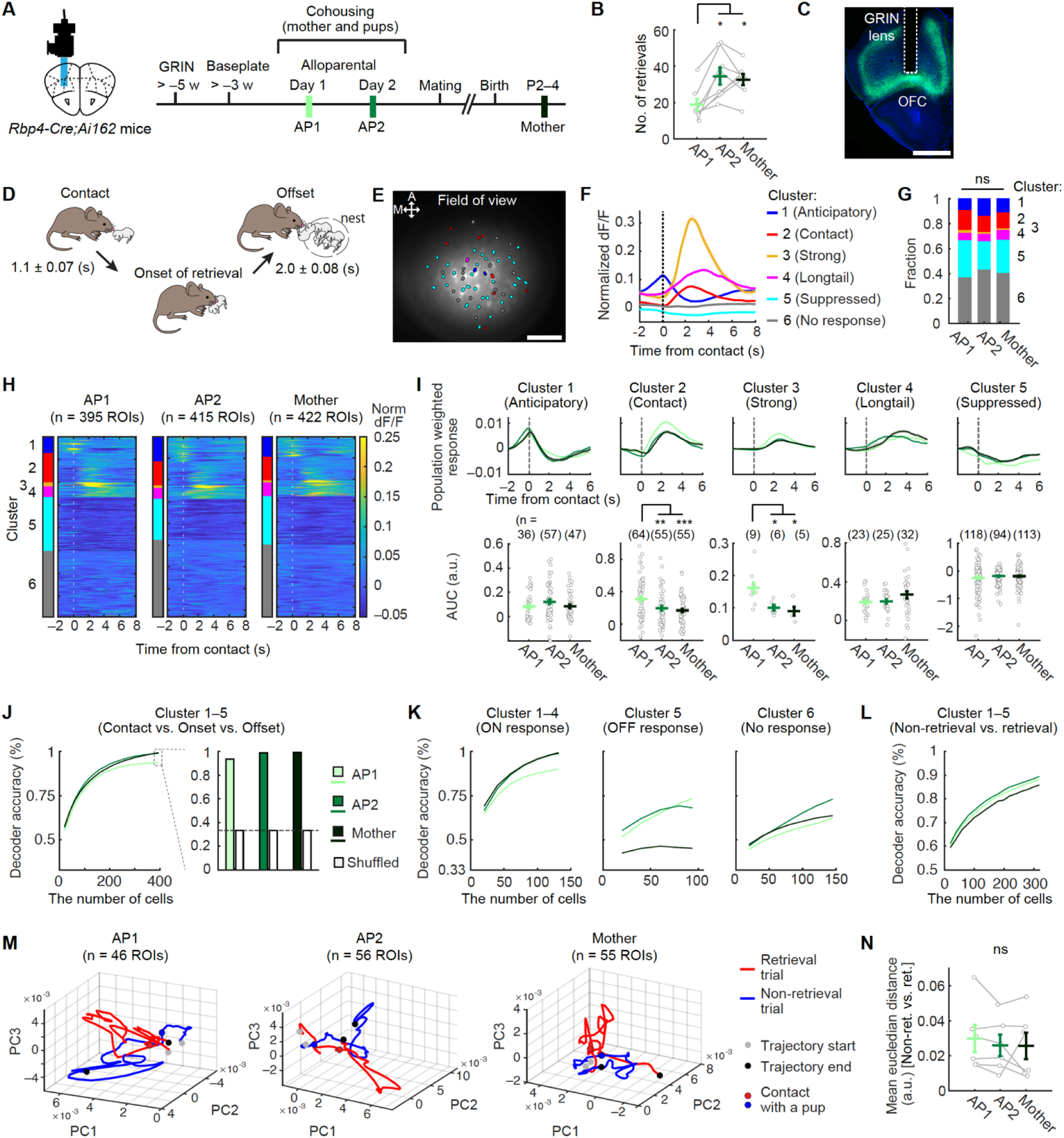
Representation of pup retrieval in OFC^Rbp4^ neurons across behavioral acquisition. (A) Experimental paradigm schematic. AP, alloparental day; P2–4, postpartum days 2–4. (B) Total retrievals during two 6-min imaging sessions. *, p < 0.05 by post hoc Tukey’s honestly significant difference (HSD) test following a significant one-way ANOVA. N = 8 mice. (C) Representative image showing the GRIN lens tract and GCaMP6s expression. Scale bar, 1 mm. (D) Schematic of sequential retrieval behaviors, indicating the average duration (±SEM) in females after 2 days of cohousing. (E) Spatial map of retrieval-responsive ROIs in a female mouse on AP2. ROIs are outlined in black, with colors indicating clusters. A, anterior; M, medial. Scale bar, 100 μm. (F) Trial-averaged normalized (norm) dF/F traces of ROIs by cluster. dF/F values were normalized to each ROI’s individual maximum. (G) Fraction of cells in each cluster. (H) Heat maps showing normalized trial-averaged responses during retrieval, sorted by cluster. Time 0 denotes pup contact followed by retrieval. ns, not significant by chi-square test with Bonferroni correction. (I) Top: population-weighted activity for each cluster. Bottom: area under the curve (AUC). *, p < 0.05, **, p < 0.01, and ***, p < 0.001 by post hoc Tukey’s HSD test after a significant one-way ANOVA. Number of ROIs shown in panel. (J) SVM decoding accuracy for classifying contact, onset, and offset of retrieval. (Left) Accuracy vs. cells used. (Right) Accuracy using 395 cells. Dotted line indicates chance. (K) Same as (J, left), using ROIs from specific clusters. (L) SVM decoding of retrieval vs. non-retrieval trials. (M) PCA trajectories from a representative mouse during 8-sec retrieval and non-retrieval epochs. (N) Mean cumulative Euclidean distance between PCA trajectories. ns, not significant by one-way ANOVA. N = 6 mice. Error bars, SEM. See Figs. S2–S5 for more data.

To monitor potential modulations in OFC^Rbp4^ neuron responses during pup retrieval across the behavioral acquisition process, we conducted chronic imaging on the same animals at various stages: before cohousing, 1 and 2 days after cohousing, and after the transition to biological motherhood (Fig. 2A). At each stage, we conducted two consecutive 6-min recording sessions in which new pups were promptly presented upon the retrieval of existing ones. To compare OFC^Rbp4^ neuron activity patterns among virgin females with different levels of retrieval performance, we designated the first day on which virgin females retrieved ten or more pups within a session as alloparental day 1 (AP1). During AP1, the total number of pup retrievals was small, and the fraction of pup contact followed by retrieval was low (Fig. 2B, S2B). By contrast, on the subsequent day of cohousing (AP2), these parameters significantly increased, and virgin females exhibited pup retrieval proficiency comparable to lactating mothers (Mother; corresponding to 2–4 postpartum days). Fig. 2C–E illustrates a representative example of the AP2 female during the pup retrieval assay, with 60 regions of interest (ROIs; putatively corresponding to individual cells) within the field of view. Pup retrieval involves three major behavioral categories (in the following sequence): pup contact, retrieval onset, and placing a pup into the nest (completion) (Fig. 2D).

To classify the heterogeneous activity patterns of OFC^Rbp4^ neurons across AP1, AP2, and Mother stages (Fig. S2C), we conducted an unbiased clustering analysis on event-averaged Ca^2+^ responses during pup retrieval (Fig. S2C–E), which revealed five distinct response clusters and one non-responsive cluster (Fig. 2F). These clusters were qualitatively labeled as follows: anticipatory (Cluster 1), contact (Cluster 2), strong (Cluster 3), longtail (Cluster 4), suppressed (Cluster 5), and non-responsive (Cluster 6). ROIs belonging to each cluster were spatially dispersed throughout the imaging field, with no evident topographical organization (Fig. S2F, G). Notably, the proportion of cells in each cluster did not significantly differ among AP1, AP2, and Mother stages (Fig. 2G). To visualize the temporal dynamics of Ca^2+^ responses, ROIs were sorted by cluster and aligned to the onset of pup contact to construct activity heat maps (Fig. 2H). This analysis shows that OFC^Rbp4^ neurons as a population represent the entire temporal structure of pup retrieval, including anticipatory periods.

To quantify stage differences, we calculated population-weighted responses by multiplying the average response of each cluster by its relative proportion. Clusters 2 and 3 showed a significant decrease in activity from AP1 to AP2 and Mother, while activity of other clusters remained unchanged (Fig. 2I). To assess the stability of individual neuron responses over the course of behavioral acquisition, we longitudinally tracked Ca^2+^ activity in a subset of ROIs between AP1 and AP2. Most ROIs retained consistent response profiles, except for Cluster 2, which showed a reduced response (Fig. S3), mirroring the trend in Fig. 2I. These data suggest that although the overall response profiles are largely stable, specific subsets of OFC^Rbp4^ neurons become selectively tuned during the acquisition of alloparental retrieval behavior.

To explore whether OFC^Rbp4^ neurons are engaged during pup retrieval even in the absence of prior pup contact, we artificially induced pup retrieval in virgin females by chemogenetically activating oxytocin (OT) neurons in the paraventricular hypothalamic nucleus (PVH) (Fig. S4A, B) (*8*). An AAV carrying *hM3D(Gq)-Myc* under the control of the *OT* promoter (*48*) was injected into the PVH, and Gq-Myc expression was confirmed via immunostaining (Fig. S4C). Administration of clozapine-N-oxide (CNO) significantly facilitated pup retrieval (Fig. S4D). Virgin females with activated OT neuron displayed robust OFC^Rbp4^ neuron activity patterns corresponding to cluster 1, 2, 5, and 6, closely resembling those observed in cohoused females (Fig. 2), although Clusters 3 and 4 were sparsely represented (Fig. S4E, F). Of note, OT neuron activation also increased the baseline activity of OFC^Rbp4^ neurons, which remained elevated on the day following CNO administration compared to baseline, suggesting a lasting modulatory effect on OFC activity (Fig. S4G). These data align with the view that OFC^Rbp4^ neurons can be rapidly recruited to respond during pup retrieval, even without prior cohousing.

### Decoding capability of OFC^Rbp4^ neurons

We next examined the decoding capability of OFC^Rbp4^ neurons across AP1, AP2, and Mother stages. We trained a linear support vector machine (SVM) to classify different behavioral categories during pup retrieval (Methods). Classification accuracy increased as more ROIs were included in the training set. Notably, decoding accuracy was higher in the AP2 and Mother stages compared to AP1, despite reduced activity in Clusters 2 and 3 in AP2 and Mother stages (Fig. 2J). In contrast, shuffling the dataset markedly impaired classification accuracy to near-chance levels. To identify which neural response types contributed to decoding, we trained separate SVMs using Clusters 1– 4 (designated as retrieval-ON), Cluster 5 (retrieval-OFF), and Cluster 6 (non-responsive). Retrieval-ON cells yielded high decoding accuracy, whereas retrieval-OFF and non-responsive cells contributed minimally (Fig. 2K). These results indicate that behaviorally responsive OFC^Rbp4^ neurons encode sufficient information to reliably distinguish behavioral categories during pup retrieval.

Considering that not all pup contact trials resulted in retrieval (Fig. S2B), we also analyzed OFC^Rbp4^ neuron activity during non-retrieval trials. Activity heat maps revealed that responses across all retrieval-ON and -OFF clusters were markedly weaker in non-retrieval trials (Fig. S5A). Quantitative analysis confirmed that response amplitudes across all these clusters were significantly lower compared to retrieval trials, despite similar pup sniffing durations across conditions (Fig. S5B, C). Longitudinal tracking of Ca^2+^ response in a subset of neurons from a naïve day to AP1 showed consistently low responses during non-retrieval trials, further supporting the idea that OFC activity is selectively enhanced during successful pup retrieval (Fig. S5D, E). We then assessed whether OFC activity during the contact period could predict retrieval outcome using an SVM. Across AP1, AP2, and Mother stages, population activity was sufficient to predict whether a trial would result in pup retrieval (Fig. 2L), supporting that OFC^Rbp4^ neurons maintain robust behavioral coding across stages.

To further explore population-level neural dynamics during pup retrieval, we generated response vectors from simultaneously recorded neurons in each mouse and visualized them using principal component analysis (PCA). We plotted the population responses during the first retrieval and first non-retrieval trial at each stage in a representative mouse (Fig. 2M). To quantify stage differences, we calculated the Euclidean distance between averaged PCA trajectories for retrieval and non-retrieval trials. Consistent with comparable decoding performance, the trajectory separation remained stable across AP1, AP2, and Mother stages (Fig. 2N). Together with the observed reduction in Clusters 2 and 3 responses in AP2 and Mother (Fig. 2I), these results suggest that while overall behavioral coding in the OFC is maintained, subtle refinements in activity patterns may contribute to enhanced retrieval performance over time.

### Substantial overlap between OFC^Rbp4^ neurons responsive to pup retrieval and reward

Mothers are innately motivated to care for their pups with strong rewards (*30*). Therefore, we hypothesized that retrieval-responsive neurons may encode aspects of positive valence. To test this, we examined the extent of overlap between OFC^Rbp4^ neurons responding to pup retrieval and those activated by a nonsocial reward. As a nonsocial reward, we used 10% sucrose solution in water-deprived mice (Fig. 3A). To assess responses to both pup retrieval and sugar water reward within the same ROIs, we conducted Ca^2+^ imaging during both pup retrieval and water licking tasks in freely moving animals (Fig. 3B, C). We identified 31 ROIs with increased responses (“Water-ON”) and 114 ROIs with suppressed responses (“Water-OFF”) in response to sugar water (Fig. 3D, left), as well as pup retrieval-responsive ROIs from the same session, classified according to clusters defined in Fig. 2 (Fig. 3D, right). Notably, 35% of Water-ON ROIs overlapped with pup retrieval-responsive Clusters 2 neurons (Fig. 3E), a proportion significantly higher than expected by chance (Fig. 3F). Overlaps with Clusters 3 and 5, however, were not significant (Fig. 3F). In contrast, Water-OFF ROIs showed significant overlap with Clusters 1 and 4 (Fig. 3G, H). While the precise functional roles of Water-ON and Water-OFF cells remain elusive, these observations support the idea that distinct pup retrieval-ON clusters may contribute differentially to reward coding (*29*).

**Fig. 3.**
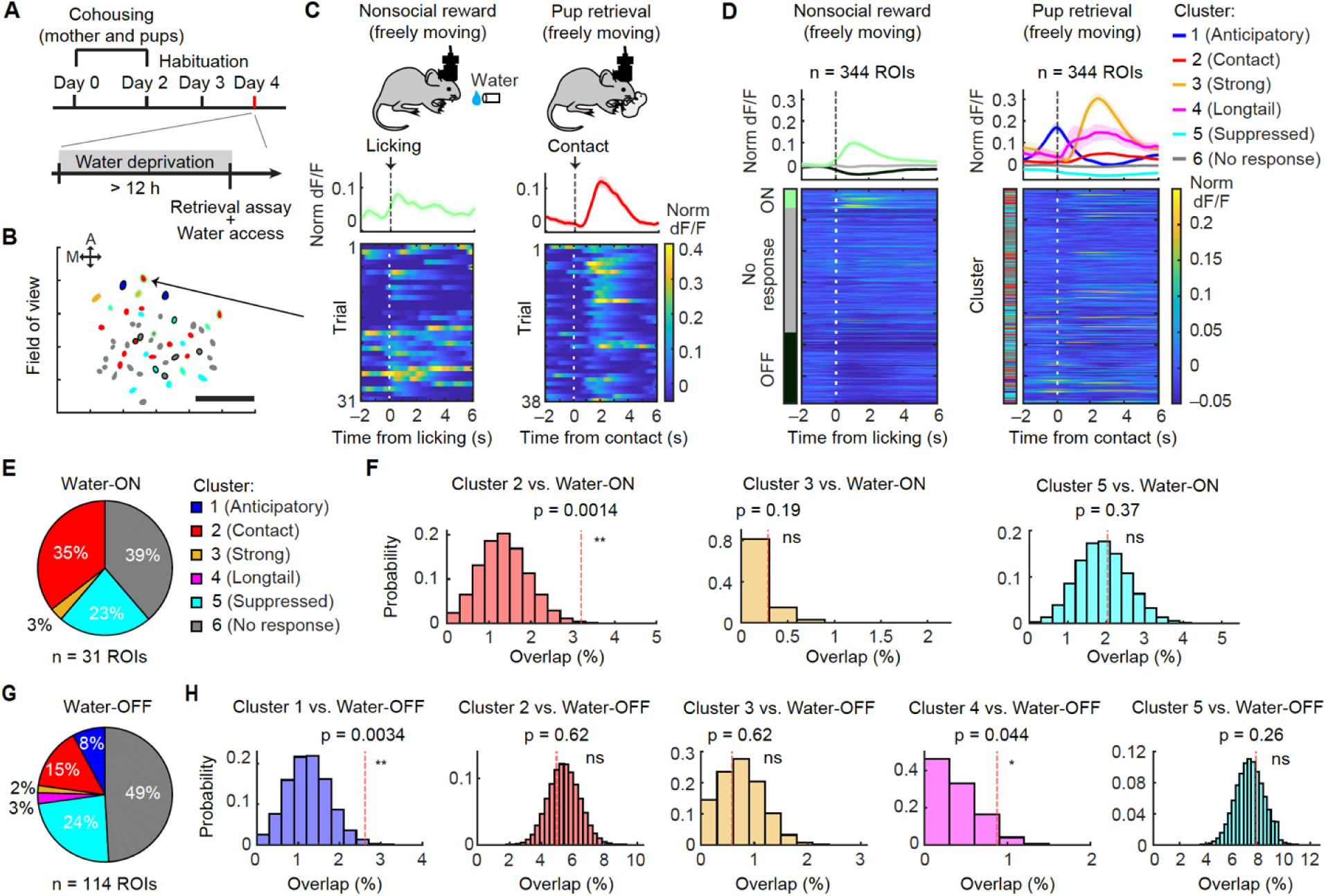
Substantial overlap between OFC^Rbp4^ neurons responsive to pup retrieval and sucrose water. (A) Schematic of the experimental paradigm. (B) Example of a spatial map of ROIs that responded during pup retrieval and passive exposure of 10% sucrose water as a nonsocial reward. A, anterior, M, medial. Scale bar, 100 μm. (C) Trial-averaged activities (top) and corresponding activity heat maps of individual trials (bottom) for a ROI responding to both retrieval and the water reward. (D) (Top) Trial-averaged activity traces of ROIs belonging to each of the three clusters during water licking (left) and each of the six clusters during pup retrieval (right). (Bottom) Heat maps showing normalized, averaged responses of individual ROIs during licking water (left) and pup retrieval (right). ROIs are sorted by their responsiveness to water licking. Time 0 indicates the moment of the lick or pup contact followed by retrieval (n = 344 ROIs from N = 6 mice). (E, G) Fraction of cells in each cluster. (F, H) Observed overlaps compared with the null distribution assuming independence between nonsocial reward-and retrieval-responsive ROIs (*, p < 0.05, and **, p < 0.01 by extreme upper-tail probability from a binomial distribution).

### OFC facilitates pup retrieval-related activities of DA neurons across multiple temporal scales

Our study thus far has revealed that OFC^Rbp4^ neurons exhibit a neural representation of pup retrieval and are required for the efficient acquisition of maternal behaviors. How do OFC^Rbp4^ neurons facilitate behavioral acquisition in the downstream neural circuit? Given that VTA^DA^ neurons can be directly regulated by OFC (*36, 38*) and critically guide maternal motivation toward infants (*33*), we next aimed to characterize VTA^DA^ neurons as a potential output of the OFC. We first confirmed the direct projections from OFC^Rbp4^ neurons to the VTA (Fig. S6A, B), as well as the presence of OFC^Rbp4^ neurons that were retrogradely labeled from the VTA, although this represents a small percentage of the population (Fig. S6C–E). Subsequently, we utilized optogenetic silencing of the OFC in combination with Ca^2+^ imaging of VTA^DA^ neurons using fiber photometry (Fig. 4A). We injected AAVs encoding the inhibitory opsin GtACR2, with YFP as a control, into the OFC of *DAT-Cre* mice (*49*) crossed with *Ai162* mice (Fig. 4B, C). Two weeks later, we bilaterally implanted optic fibers just above the OFC for optogenetic manipulation and another one into the left VTA for fiber photometry (Fig. S6F, G). We validated that GCaMP6s was selectively expressed by DA neurons in the *DAT-Cre*; *Ai162* mice using fluorescent *in situ* hybridization (Fig. 4D).

**Fig. 4.**
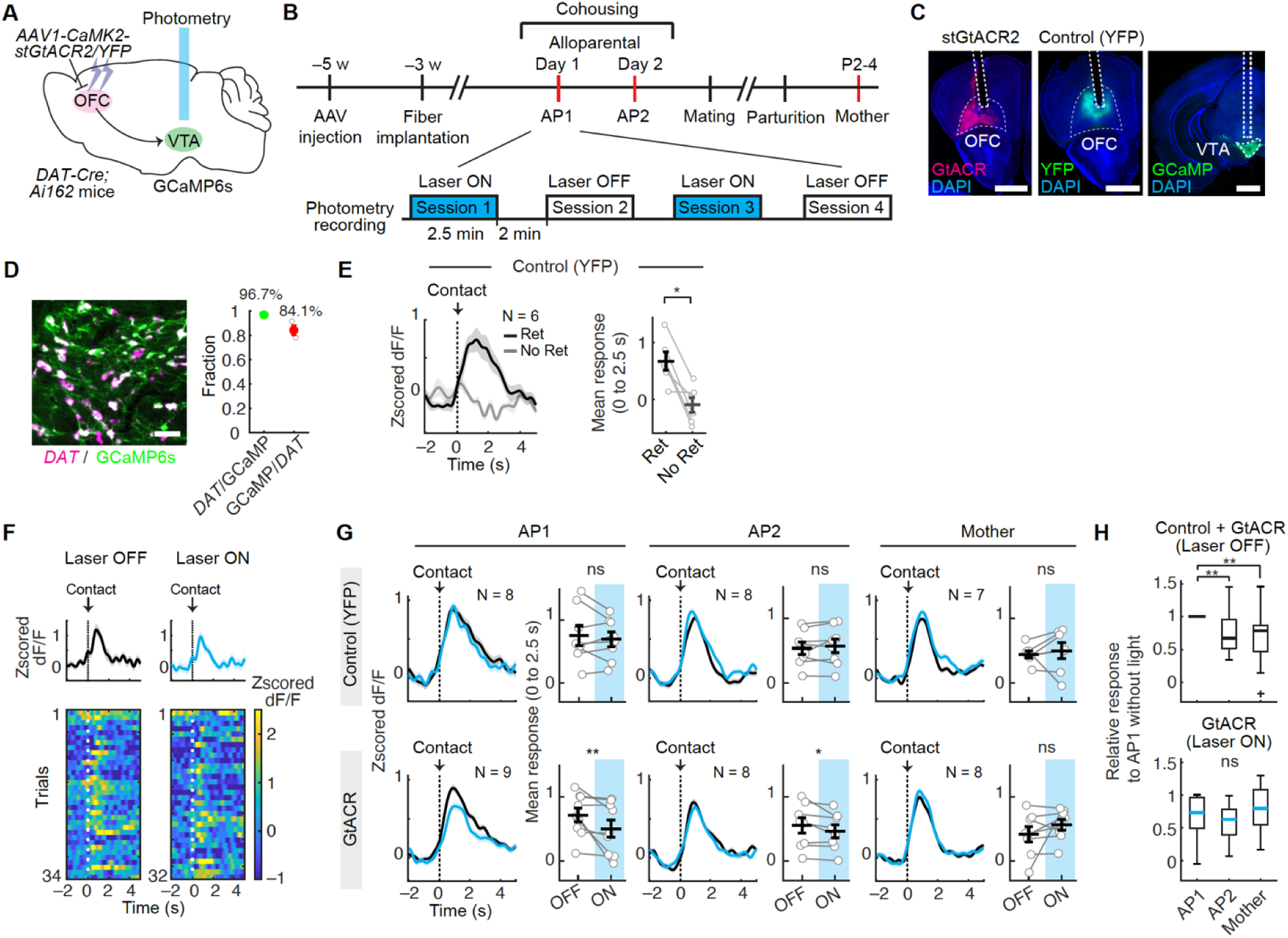
The OFC facilitates pup retrieval-related activities of VTA^DA^ neurons. (A) Schematic of the experimental design. AAV1 *CaMKII-stGtACR2-fusionRed* or AAV1 *CaMKII-YFP* was injected into the bilateral OFC of virgin *DAT-Cre; Ai162* mice. (B) Experimental timeline: four imaging sessions (2.5 min each); first/third with laser (blue), second/fourth as internal controls without laser. (C) Representative coronal sections showing the optic fiber tract and expression of stGtACR2, YFP, and GCaMP6s in *DAT-Cre; Ai162* mice. Scale bar, 1 mm. (D) (Left) Representative coronal section showing *DAT* mRNA (magenta) and GCaMP6s (anti-GFP, green) in *DAT-Cre; Ai162* mice. (Right) Quantification of specificity (*DAT*/GCaMP6s) and efficiency (GCaMP6s/*DAT*) (N = 3 mice). Scale bar, 50 μm. (E) (Left) Trial-averaged Z-scored peri-event time histograms (PETHs) of control animals during contact followed by retrieval (Ret, black line) and non-retrieval (Not ret, gray line). Shadow represents the SEM. (Right) The mean of Z-scored PETHs between 0 and 2.5 s, aligned to pup contact. *, p < 0.05 by Wilcoxon signed-rank test. (F) Trial-averaged activities (top) and heat maps (bottom) of VTA^DA^ neurons during pup retrieval with and without OFC inactivation in an AP2 mouse. Time 0 indicates pup contact followed by retrieval. (G) Z-scored PETHs with (blue) and without (black) laser, and mean Z-scored PETHs (0–2.5 s). (Top) YFP-injected group. (Bottom) GtACR2-injected group. The number of animals (N) is indicated in the panel. *, p < 0.05, and **, p < 0.01 by Wilcoxon signed-rank test. ns, not significant. (H) Normalized mean response for pup retrieval across the AP1, AP2, and Mother stages. Laser-off data pooled from GtACR (N = 9 mice) and YFP (N = 8 mice) groups as control. ns, not significant, and **, p < 0.05 by a post hoc Tukey’s HSD test following a significant Kruskal–Wallis test. See Fig. S6 for more data.

We conducted four sessions of pup retrieval assays, each lasting 2.5 min, while simultaneously performing imaging, with and without laser stimulation (Fig. 4B). Throughout each session, new pups were presented immediately upon the retrieval of existing ones. To assess the influence of silencing OFC neurons on VTA^DA^ neuron activity, we compared the Ca^2+^ transients of VTA^DA^ neurons with and without laser stimulation around the peri-events of pup retrievals. Consistent with previous reports (*11, 33*), VTA^DA^ neurons exhibited time-locked activity following pup contact (Fig. 4E). Non-retrieval trials did not show a significant photometry signal of VTA^DA^ neurons, indicating that the phasic activity of these neurons during pup retrieval is behaviorally relevant. Silencing the OFC led to a significant decrease in the Ca^2+^ activity of VTA^DA^ neurons in the AP1 and AP2 stages (Fig. 4F, G), without affecting general locomotor activity (Fig. S6H). The reduction in VTA^DA^ neuron activity was more pronounced in AP1 than in AP2, while no such effect was observed in the Mother stage. Taken together, these data show that the OFC facilitates VTA^DA^ neuron activity during pup retrieval, particularly in the early stages of maternal behavioral acquisition.

Consistent with recent research proposing that VTA^DA^ neurons encode RPE during retrieval learning (*33*), we also observed a significant overall decline in retrieval-related activities of VTA^DA^ neurons as animals progressed from AP1 to AP2 (Fig. 4H, top). By contrast, silencing of the OFC attenuated this reduction (Fig. 4H, bottom), suggesting that the OFC is involved in the experience-dependent plasticity of VTA^DA^ neuron activity. Given that OFC^Rbp4^ neurons in Clusters 2 and 3 also exhibited reduced activity in AP2 and Mother compared to AP1 stage (Fig. 2H), changes in OFC^Rbp4^ neuron activity may underlie the performance-dependent reduction of VTA^DA^ neuron activity.

We next investigated the potential role of the OFC in modulating the temporal dynamics of VTA^DA^ neurons in a trial-by-trial manner (Fig. 5A). In the absence of OFC inactivation, we observed a more intensive response during the first five as compared to the last five retrieval trials, as inferred by the peak height of the photometric signals, across AP1, AP2, and Mother (Fig. 5B). By contrast, the silencing the OFC canceled the differences between the first and last five trials in AP1 and AP2, while the OFC no longer affected the dynamics of VTA^DA^ neuron activity within the session in the Mother stage. These data indicate that i) VTA^DA^ neuron activity rapidly adapts during individual sessions, even within a single day, a phenomenon we term the “adaptation”, and ii) the OFC is involved in encoding the adaptation in virgin females, while its influence on the adaptation disappears during the biological transition to motherhood.

**Fig. 5.**
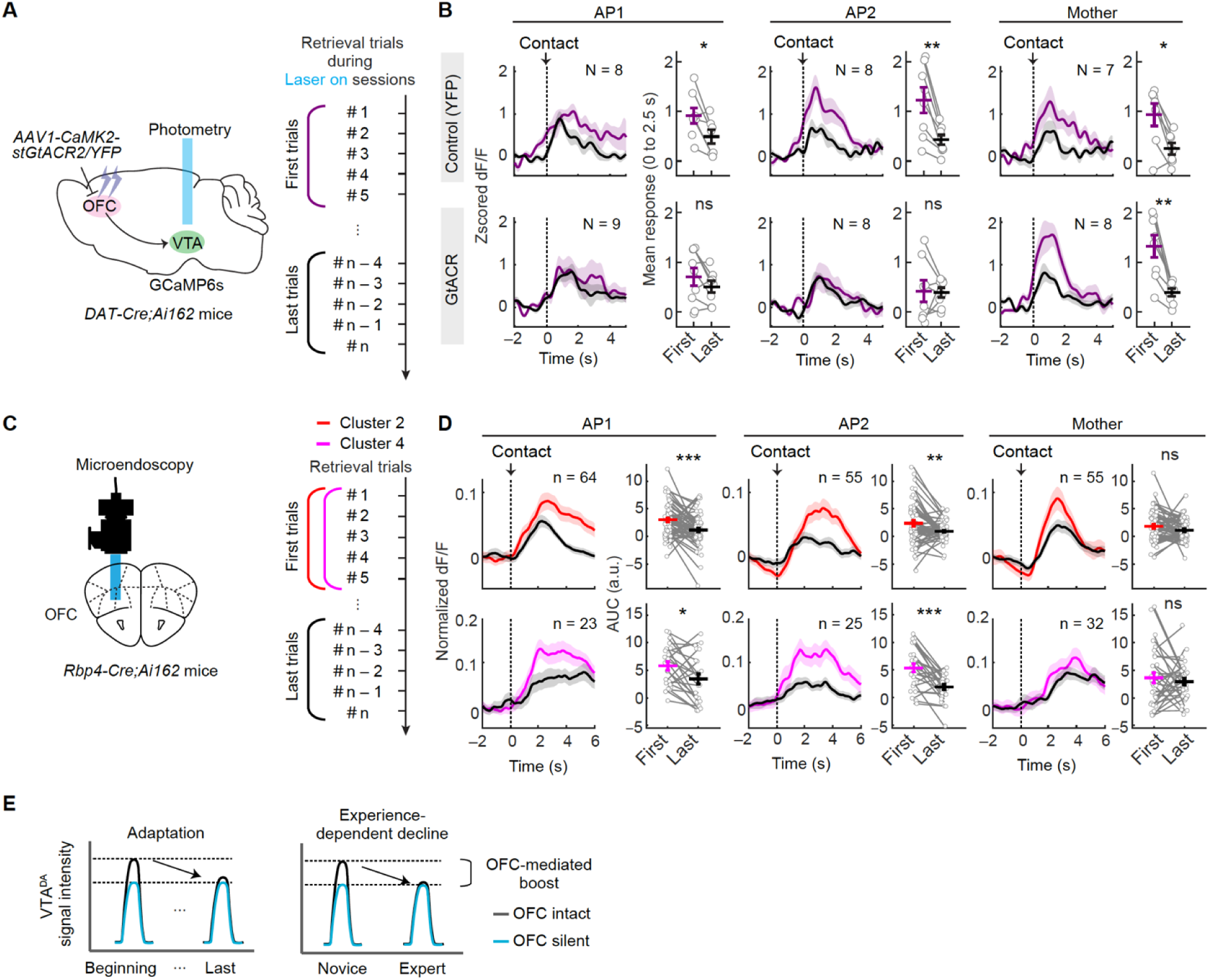
The OFC regulates the dynamic modulation of VTA^DA^ neuron activity. (A, C) Schematics of the experimental design and analyzed trials. (B) Z-scored PETHs for the first (black) and last (purple) five trials during laser-on sessions for the control (top) and GtACR2 (bottom) groups. The right graph represents the mean of Z-scored PETHs from 0 to 2.5 s, aligned to pup contact. *, p < 0.05, and **, p < 0.01 by Wilcoxon signed-rank test. The number of animals (N) is indicated in the panel. (D) Averaged activity traces of ROIs from Cluster 2 (top) and Cluster 4 (bottom) for the first (black) and last (purple) five pup retrieval trials. The right graph shows the AUC of normalized dF/F (0–4 s for Cluster 2; 0–6 s for Cluster 4) aligned to pup contact. *, p < 0.05, and ***, p < 0.001 by paired t-test. The number of ROIs is indicated in the panel. (E) Schematic describing two models of VTA^DA^ neuron dynamics and the contribution of the OFC. Shadow represents SEM. Time 0 indicates pup contact followed by retrieval. See Fig. S7 for more data.

We further examined potential adaptation in the microendoscopic OFC imaging data presented in Fig.2 (Fig. 5C). Clusters 2 and 4 exhibited within-session adaptation patterns resembling those observed in VTA^DA^ neurons, particularly during the AP1 and AP2 stages (Fig. 5D). This is consistent with our finding that OFC silencing attenuated adaptation effects in VTA^DA^ neurons specifically during the AP1 and AP2 stages, but not during the Mother stage (Fig. 5B). In contrast, no such adaptation was detected in the remaining clusters (Fig. S7). These results raise the possibility that specific subsets of OFC^Rbp4^ neurons may contribute to the modulation of VTA^DA^ neuron activity, although direct causal relationships remain to be elucidated in future studies.

### OFC facilitates DA release into the ventral striatum (VS)

VTA^DA^ neurons send their axons to various downstream regions, including different subregions of the striatum and amygdala (*50*). Although a recent study utilizing a genetically encoded fluorescent DA sensor (*51*) demonstrated DA release into the VS during pup retrieval (*52*), the regional diversity of DA signaling during pup retrieval remains unexplored. To address this issue, we virally targeted GRAB^DA3m^ to the following four specific regions for photometric recording: the VS, dorsal striatum (DS), posterior dorsolateral striatum (pDLS), and basolateral amygdala (BLA) (Fig. 6A, B). Representative traces of regional DA signals are shown in Fig. 6C. We observed significant DA transients during pup retrieval compared with the non-retrieval trials across all regions, suggesting their potential contribution to pup retrieval behaviors (Fig. 6D). Specifically, area under the receiver operating characteristic curve (AUC-ROC) analysis showed that the DA transients reached statistical significance just before or after pup contact across all the regions (Fig. 6E). To assess the regional diversity of DA signaling, we compared the phasic DA traces among the different regions (Fig. 6F). We observed the largest DA transients during the pup retrieval trials in the VS (Fig. 6F), with the BLA exhibiting a longer latency to peak than the other regions. Furthermore, the VS exhibited a larger full-width-half-maximum (FWHM) than the DS and pDLS, indicating the region-specific temporal dynamics of DA transients. These findings highlight the heterogeneous nature of DA signaling and its temporal regulation across targeted subregions during pup retrieval.

**Fig. 6.**
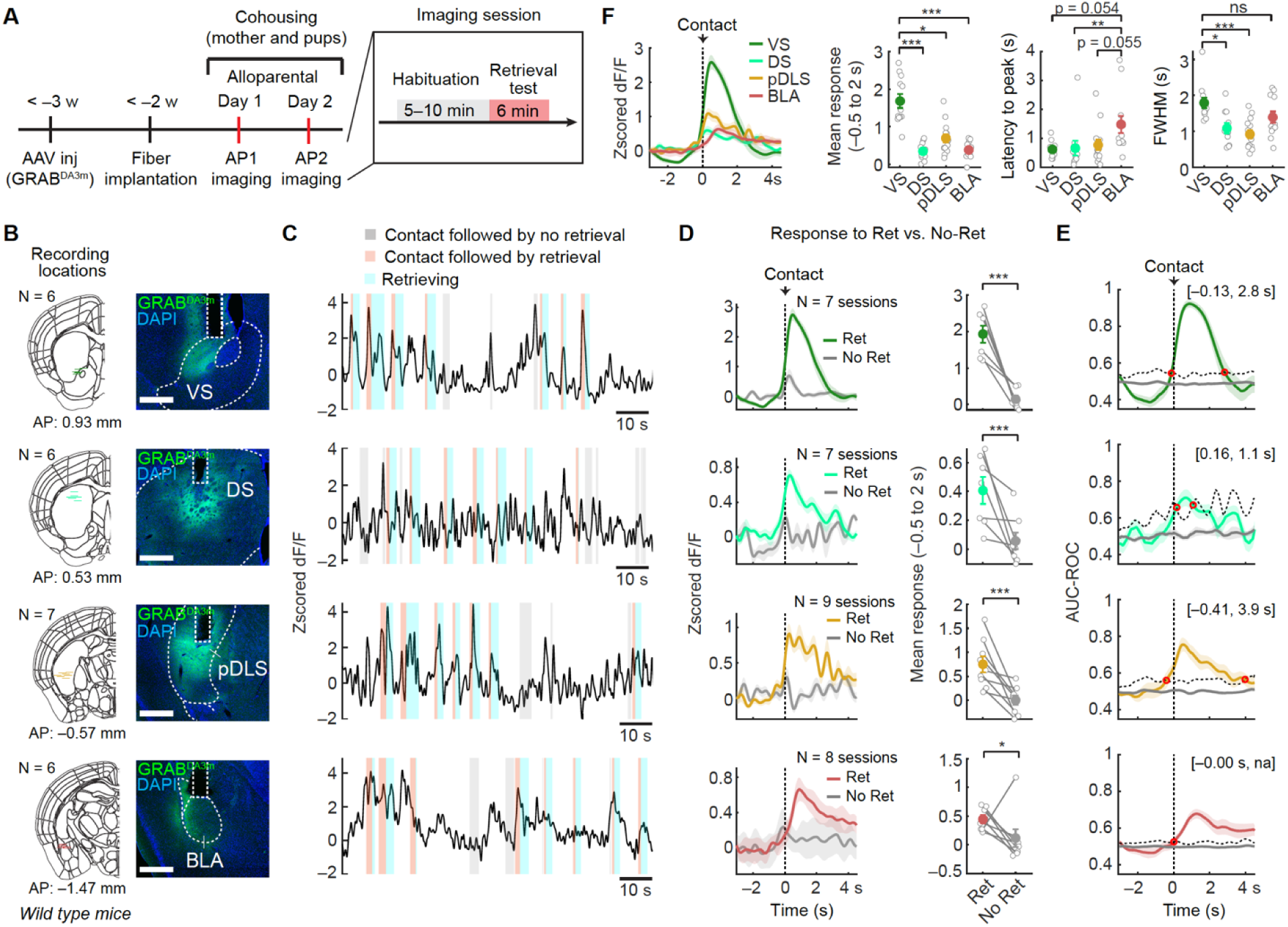
DA is released in the multiple downstream targets during pup retrieval. (A) Schematic of the experimental timeline. (B) (Left) Schematics of coronal sections showing the position of the optic fiber (N = 6–7 mice each). AAV9 *hSyn-GRAB^DA3m^* was injected into the left VS, DS, pDLS, or BLA of the virgin wild-type mice. (Right) Representative coronal sections showing the expression of GRAB^DA3m^ without antibody staining. Scale bar, 500 μm. (C) Representative photometry trace recorded from an AP2 female during pup retrieval. Colors correspond to specific behavioral events as indicated above the panel. (D) (Left) Trial-averaged Z-scored PETH traces fitted to pup contact followed by retrieval (colored) or non-retrieval (gray). (Right) The mean of Z-scored PETHs between –0.5 and 2 s, aligned to pup contact. The number of sessions is indicated in the panel (data pooled from AP1 and AP2). *, p < 0.05, and ***, p < 0.001 by the Mann–Whitney *U* test. (E) The averaged traces of the AUC-ROC calculated for each animal. Dashed black lines display the mean +2 standard deviation (SD) of the AUC-ROC from trial-shuffled data (gray trace). Red dots indicate time points exceeding or falling below 2SD of shuffled data for the first time.

Lastly, we investigated the role of the OFC in DA release in the VS, given the most pronounced DA transient observed, along with previous research suggesting the importance of the VTA–VS circuit in maternal behaviors (*53, 54*). Due to spatial constraints, optogenetic manipulation in the OFC influenced GRAB^DA3m^ signals in the VS. Hence, we utilized a chemogenetic approach. An AAV expressing Cre-dependent hM4Di(Gi)-mCherry was injected into the bilateral OFC of *Rbp4-Cre* animals, along with another AAV expressing GRAB^DA3m^ into the VS (Fig. 7A). We monitored the GRAB^DA3m^ signal during pup retrieval while the OFC^Rbp4^ neurons were silenced by CNO administration (Fig. 7B). Post hoc histological analyses confirmed the precision of targeting for hM4Di-mCherry and the GRAB sensor (Fig. 7C, D). While CNO administration in mCherry+ control animals showed no impact on DA release during pup retrieval, chemogenetic silencing of OFC^Rbp4^ neurons resulted in a significant reduction in DA release in the VS in AP1, but not in AP2 or Mother (Fig. 7E). Of note, CNO administration did not affect locomotor activity in either the control or experimental groups (Fig. S8). These data align with the findings that OFC silencing has a more pronounced impact on VTA^DA^ neuron activity in AP1 (Fig. 4G). Collectively, these data demonstrate that the OFC facilitates DA release into the VS during pup retrieval, particularly in the early stages of maternal behavioral acquisition, potentially contributing to the efficient acquisition of maternal behaviors.

**Fig. 7.**
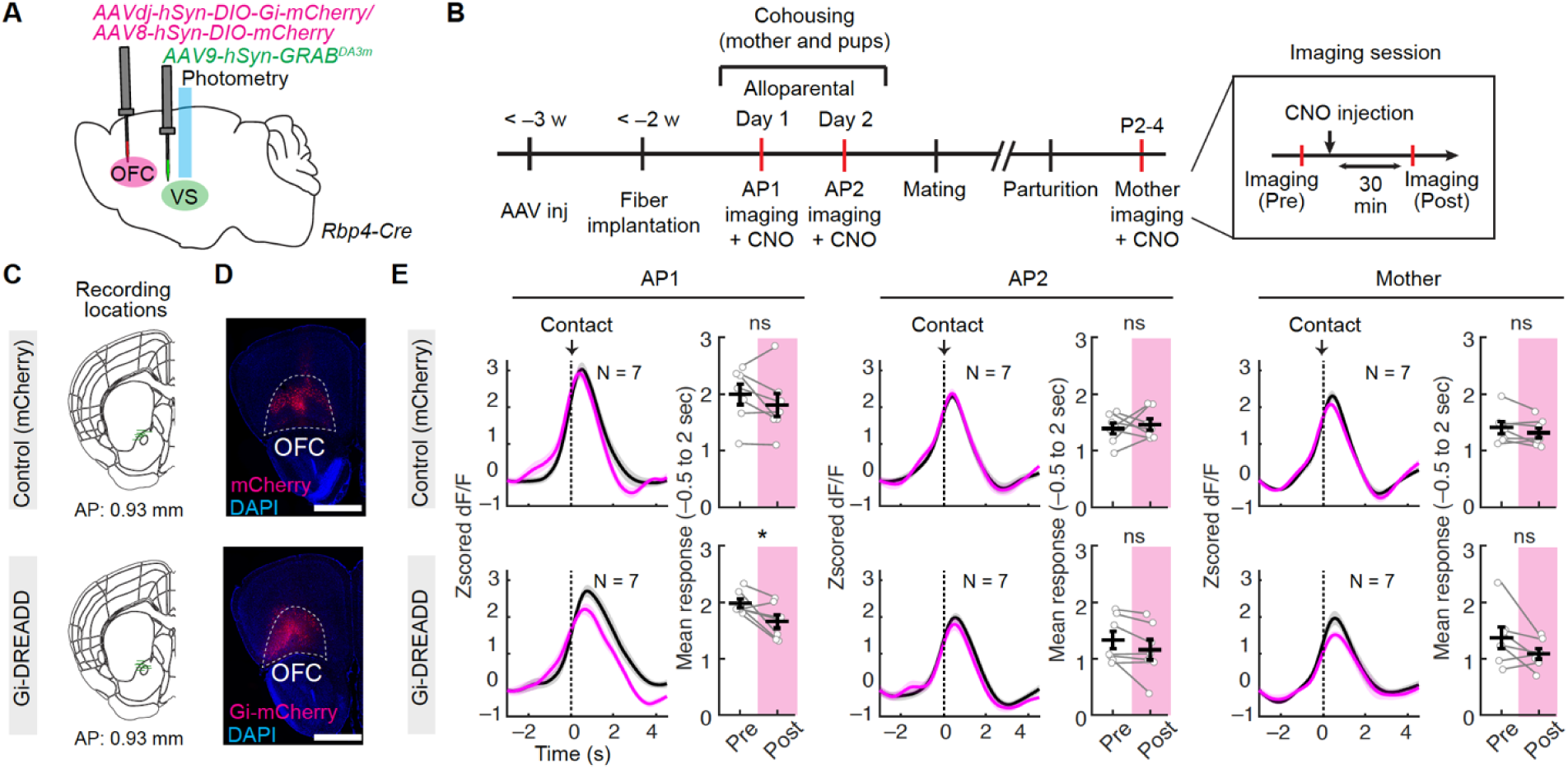
The OFC regulates DA release in the VS during pup retrieval. (A) Schematic of the experimental design. AAVdj *hSyn-DIO-Gi-mCherry* or AAV8 *hSyn-DIO-mCherry* as a control was injected into the bilateral OFC of the virgin *Rbp4-Cre* mice. AAV9 *hSyn-GRAB^DA3m^* was injected into the left VS. (B) Schematic of the experimental time line, including an imaging session (duration: 6 min). (C) Schematics of coronal sections showing the position of the optic fiber (N = 7 mice for each group). (D) Representative coronal sections showing the expression of mCherry (top) and Gi-mCherry (bottom). Scale bar, 1 mm. (E) Z-scored PETHs before (black line) and after (magenta line) CNO administration (left), with corresponding mean responses of Z-scored PETHs. (right). Top: mCherry-injected control group. Bottom: hM4Di-injected group. The number of animals is indicated in the panel. Time 0 indicates pup contact. *, p < 0.05, by the Wilcoxon signed-rank test. ns, not significant. See Fig. S8 for more data, and S9 for schematic summary of all the data.

## Discussion

During the last decade, there has been intensive research on the limbic circuits for parental behaviors (*10, 55*). However, our understanding of the potential functions of PFC in parental behaviors remains limited. Given the substantial evolutionary expansion of the human frontal cortex, understanding the higher-order functions associated with parental behaviors and the potential interactions between the frontal cortex and limbic circuits during such behaviors could be expected to offer valuable insights into human parental behaviors. The present study characterizes the functions of the OFC that modulates the plasticity of the DA system (Fig. S9), although it remains unclear whether this regulation is mediated by direct projections from OFC^Rbp4^ neurons to VTA^DA^ neurons or via indirect, multi-synaptic pathways. Here, we discuss the insights obtained from our study and its limitations.

### Roles of the OFC in maternal behaviors

In contrast to the parental behavior centers in the preoptic area, such as galanin neurons (*1, 10*) or Calcr neurons (*13*), whose dysfunction severely blocks maternal caregiving, our data suggest that the OFC has a modulatory role on the acquisition of parental behaviors. Both cell ablation and optogenetic inhibition of the OFC delayed the acquisition of maternal behaviors following cohousing with mothers and pups or repeated exposures to pups (Fig. 1), implying the involvement of the OFC in linking pups with innate values. Supporting this notion, OFC^Rbp4^ neurons responsive to pup retrieval also exhibited a significant overlap in response to nonsocial rewards (Fig. 3). Moreover, the OFC played a role in modulating DA neuron activity (Figs. 4 and 5) and DA release in the VS (Fig. 7), a core component of the brain’s reward system. Collectively, the OFC may enhance motivation to drive maternal behaviors during the process of behavioral acquisition, conceptualizing pups as rewards (Fig. S9).

While decades of research using sophisticated learning paradigms have shown the role of the OFC in value-based decision-making and reversal learning by representing and updating subjective values (*23, 56, 57*), relatively little attention has been paid to the neural representations of the OFC during naturally occurring behavioral acquisition. Our data offer several insights. First, the OFC harbors a largely stable representation of pup retrieval (Fig. 2), which can emerge even without prior experience of pup retrieval (Fig. S4). This may reflect the intrinsic value and motor aspects of instinctive behaviors, in contrast to the prevailing view that PFC representations of social behavior are predominantly shaped experience or learning (*58-60*). Second, within this stable framework, we observed notable refinements, including a significant reduction in pup contact-related activity across days (Fig. 2) and within sessions (Fig. 5). These changes parallel the decline in activity of downstream VTA^DA^ neurons (Figs. 4 and 5). While we interpret these declines as reflecting fine-tuning and adaptation, their precise roles in pup retrieval remains to be determined in future studies.

We recognize several limitations in the present study. First, despite our focus on the OFC due to its neural activation by infant-related sensory inputs in both humans and mice (*4, 21, 22*), we did not examine regional specificity of the OFC in regulating maternal behavioral acquisition. Other areas of the frontal cortex, such as the mPFC (*15*) and the anterior cingulate cortex (*61*), may have significant roles alongside the OFC. In addition, although our study focuses on layer 5 of the OFC, given that subcortical projections primarily originate from this deep layer (*39, 62*), other layers, particularly layer 6 (see Fig. S6D), may also contribute substantially to VTA projections, warranting further investigation. Second, although we found OFC^Rbp4^ neurons whose activities tiled all behavioral categories of pup retrieval, it remains unclear whether these distinct populations are associated with subtypes of neurons based on their axonal projection or gene expression profiles (*63, 64*). As projection-defined neurons can encode unique or overlapping information (*36, 65*), future studies should address whether transcriptional types or projection-defined neurons in the OFC convey unique information about pup retrieval to specific downstream targets. Third, we acknowledge that variations in internal states may complicate comparisons of the responsiveness of OFC^Rbp4^ during pup retrieval to nonsocial rewards (Fig. 3). For instance, the reward magnitude depends on the internal state, as sugar water does not serve as a reward for a satiated animal (*66*). Future studies should develop imaging or recording protocols capable of analyzing multiple task categories where reward scales can be controlled.

### Roles of the DA system in maternal behaviors

While previous studies have established roles of VTA^DA^ neurons in maternal behaviors (*34*), the mechanisms underlying the temporal dynamics of their neural activity during pup retrieval (*11, 33*) have remained elusive. Our data suggest a functional link by which OFC neurons facilitate pup retrieval-locked VTA^DA^ neuron activity only during the early stages of behavioral acquisition (Fig. 4). However, a key limitation of our study is the absence of pathway-specific manipulations targeting the OFC-to-VTA projection, leaving the precise role of the direct pathway unresolved. As behavioral performance improves, the OFC-mediated enhancement of VTA^DA^ activity gradually disappears, which accounts for the experience-dependent gradual decline in VTA^DA^ activity (*33*) (Fig. 4). In addition, analogous to the known representation of behavioral novelty and saliency in VTA^DA^ neurons (*67*), our identified adaptation effect (Fig. 5) may play roles in learning, reinforcement, and motivation. In sensory systems, neural adaptation to repetitive stimuli often enhances sensitivity to unexpected stimuli (*68*). Thus, one possibility is that VTA^DA^ neuron adaptation enhances the detectability of novel or unexpected behavioral outcomes.

A recent study proposed a social RPE model (*33*), suggesting that the experience-dependent reduction in VTA^DA^ neuron activity. In line with this, we found that inactivation of the OFC abolished the RPE-like decline in VTA^DA^ neuron activity, raising the possibility that OFC activity contributes to the generation of social RPE signals. Specifically, subsets of OFC^Rbp4^ neurons, such as those responsive to both pup contact and non-social reward (Cluster 2; Figs. 2 and 3), may be involved in this computation. While the concept of RPE signals is instrumental in enabling animals to update internal representations of their environment and adapt their behavior, the biological implementation of RPE in maternal behaviors remains largely unknown. Therefore, it is important to investigate how the OFC loses its influence on VTA^DA^ neuron activity during the transition to motherhood. While this loss of influence may be linked to maternal plasticity within the direct axonal projections from the OFC to VTA (Fig. S6) (*35, 36*), indirect pathways involving GABAergic or glutamatergic neurons (*38*) could also contribute. Additionally, understanding how VTA^DA^ neuron activity is modulated in mothers after OFC involvement wanes is important. One likely candidate is the medial preoptic area, where estrogen receptor alpha+ neurons facilitate VTA^DA^ neuron activity during pup retrieval by disinhibiting VTA^DA^ neurons (*11*). Supporting a broader framework, a recent study showed that VTA^DA^ neurons are required for the acquisition, but not the expression, of male aggressive behavior, suggesting a generalizable, transient role for VTA^DA^ neuron activity in behavioral acquisition (*69*). Future studies should investigate how circuit-level plasticity within and beyond the OFC-VTA axis governs both the acquisition and maintenance of maternal behaviors.

Subtypes of DA neurons, based on their molecular profiles and anatomical positions, project their axons toward partially overlapping yet distinct downstream targets (*36, 70*). This organization contributes to variations in DA release, which may not directly correlate with overall somatic neural activity (*71, 72*). Recent rodent studies have elucidated the functional heterogeneity among VTA^DA^ neurons across various targeted regions. For instance, DA release in the VS is linked to reward, motivation, and learning, whereas in the DS, it regulates motor function (*71, 72*). DA release in the tail of the striatum is associated with processing aversive stimuli (*73*), and in the BLA, it modulates emotional processing and fear conditioning (*74*). Despite these advancements, the dynamics of DA release during maternal behaviors have only been characterized in the VS (*52*). Our data not only support pronounced DA release in the VS during pup retrieval, but also demonstrate the presence of reliable phasic DA transients with various temporal dynamics observed across all regions, including the DS, pDLS, and BLA. Consequently, we expect that future research will focus on recording VTA^DA^ neurons at single-cell resolution and manipulating their specific projection types to elucidate the heterogeneous roles played by distinct subtypes of VTA^DA^ neurons in the regulation of maternal behaviors.

## Materials and Methods

### Animals

All animals were housed under a regular 12-h dark/light cycle with *ad libitum* access to food and water. Wild-type FVB mice were purchased from CLEA Japan, Inc. (Tokyo, Japan) for backcrossing. *DAT-Cre* (Jax# 006660) and *Ai162* (*TIT2L-GC6s-ICL-tTA2*, Jax#031562) were purchased from the Jackson Laboratory. *Rbp4-Cre* mice were originally obtained from the Mutant Mouse Regional Resource Center. We used the F1 hybrid of C57BL/6 and FVB strain for all experiments. *Rbp4-Cre* heterozygous female mice (age range, 2–5 months) were used for ablation experiments using taCasp3, optogenetic silencing experiments using GtACR2, and chemogenetic silencing experiments with photometry recording of DA dynamics using GRAB^DA^. *Rbp4-Cre; Ai162* double heterozygous female mice (age range, 2–6 months) were used for *in vivo* microendoscopic imaging of the OFC. *DAT-Cre; Ai162* double heterozygous female mice (age range, 2–6 months) were used for the fiber photometry experiments. Wild-type female mice (age range, 2–5 months) were used for the fiber photometry recordings of DA dynamics. To investigate maternal caregiving behaviors, this study exclusively utilized female mice. A similar study involving male mice will be detailed in future publications.

### Viral preparations

The following AAV vectors were generated by Addgene.

AAV serotype 1 *EF1a-FLEx-taCasp3-TEVp* (5.8 × 10^12^ gp/mL) (Addgene #45580)(*45*)

AAV serotype 5 *EF1a-DIO-YFP* (1.3 × 10^13^ gp/mL) (Addgene #27056)

AAV serotype 8 *hSyn-DIO-mCherry* (2.2 × 10^13^ gp/mL) (Addgene #50459)

AAV serotype 1 *CaMK2a-stGtACR2-FusionRed* (1.3 × 10^13^ gp/mL) (Addgene #105669) (*75*)

AAV serotype 1 *hSyn-SIO-stGtACR2-FusionRed* (1.9 × 10^13^ gp/mL) (Addgene #105677) (*75*)

AAV serotype 1 *CaMK2a-EYFP* (1.0 × 10^13^ gp/mL) (Addgene #105622)

AAV serotype retrograde *CAG-tdTomato* (1.3 × 10^13^ gp/mL) (Addgene #59462)

The following AAV vectors were generated by the UNC viral core.

AAV serotype 5 *OTp-hM3Dq-Myc* (2.9 × 10^12^ gp/mL) (Corresponding plasmid: Addgene #184753) (*48*)

AAV serotype 9 *hSyn-FLEx(FRT)-mGFP* (7.4 × 10^12^ gp/mL) (Corresponding plasmid: Addgene #71761) (*36*)

The following AAV vectors were generated by WZ Biosciences.

AAV serotype 9 *hSyn-GRAB^DA3m^*(2.2 × 10^13^ gp/mL)

*pAAV CAG-FLEx-FLPo* was constructed by in-fusion-based PCR cloning utilizing the following two DNA fragments: i) SalI-AscI restriction fragment of *pAAV CAG-FLEx-TCb* (Addgene #48332) as a vector backbone, and ii) PCR fragment of *FLPo* originated from *pTCAV-FLEx-FLPo* (#67829, Addgene). Because a nuclear localization signal (NLS) was not included in this version of the *FLPo* cassette, subsequently, we performed PCR-based subcloning of SV40 NLS to the 5’ end of the *FLPo* cassette, resulting in the construction of *pAAV CAG-FLEx-FLPo*. With this plasmid, the AAV serotype retro *CAG-FLEx-FlpO* (5.5 × 10^12^ gp/mL) was generated by the National Institute for Physiological Science viral vector core facility. The AAV serotype DJ *hSyn-DIO-hM4Di-mCherry* (6.9 × 10^12^ gp/mL) was generated by the Fukushima Medical University School of Medicine Viral Vector Core facility with the corresponding plasmid (Addgene #44362).

### Stereotaxic injection

For targeting AAV or rabies virus into a certain brain region, stereotaxic coordinates were first defined for each brain region based on the Allen Brain Atlas (*76*). Mice were anesthetized with 65 mg/kg ketamine (Daiichi-Sankyo) and 13 mg/kg xylazine (Sigma-Aldrich) via intraperitoneal injection and head-fixed to the stereotaxic equipment (RWD). For the ablation experiments (Figure 1A–G), 200 nL of AAV1 *EF1a-FLEx-taCasp3-TEVp* was injected into the right hemisphere or bilateral OFC (coordinates relative to the bregma: anteroposterior [AP] 2.5 mm, mediolateral [ML] 1.2 mm, dorsoventral from the brain surface [DV] 1.9 mm) at a speed of 50 nL/min using a UMP3 pump regulated by Micro-4 (World Precision Instruments). For the optogenetic silencing experiments (Fig. 1H–R), 200 nL of AAV1 *hSyn-SIO-stGtACR2-FusionRed* or AAV8 *hSyn-DIO-mCherry* was injected into the bilateral OFC. To activate OT neurons in the PVN (Fig. S4), 200 nL of AAV5 *OTp-hM3D(Gq)-Myc* was injected into the bilateral PVN (coordinates relative to the bregma: AP –0.5 mm, ML 0.2 mm, DV 4.4 mm). For the optogenetic experiments with photometry recordings (Fig. 4), 200 nL of AAV1 *CaMK2a-stGtACR2-FusionRed* or AAV1 *CaMK2a-EYFP* was injected into the bilateral OFC. For anterograde tracing from the OFC (Fig. S6A and B), 100 nL of a 1:1 mixture of AAVretro *CAG-FLEx-FlpO* and AAV9 *hSyn-FLEx(FRT)-mGFP* was injected into the left OFC. Of note, we utilized this AAVretro *CAG-FLEx-FlpO* just to drive FlpO into the injection site and did not intend its retrograde transductions. To calculate the proportion of OFC^Rbp4^ neurons that project to VTA (Fig. S6C–E), 100 nL of AAVretro *CAG-tdTomato* was injected into the left VTA (coordinates relative to the bregma: AP –2.7 mm, ML 0.5 mm, DV 4.4 mm). For the photometry recording of DA dynamics (Figs. 6, 7), 200 nL of AAV9 *hSyn-GRAB^DA3m^* was injected into the VS (coordinates relative to the bregma: AP 1.6 mm, ML 1.2 mm, DV 4.0 mm), DS (coordinates relative to the bregma: AP 1.0 mm, ML 1.9 mm, DV 2.7 mm), pDLS (coordinates relative to the bregma: AP –0.4 mm, ML 3.0 mm, DV 3.55 mm), or BLA (coordinates relative to the bregma: AP –0.8 mm, ML 3.1 mm, DV 4.4 mm). For the pharmacogenetic experiments with photometry recordings (Fig. 7), 200 nL of AAVdj *hSyn-DIO-hM4Di-mCherry* or AAV8 *hSyn-DIO-mCherry* was injected into the bilateral OFC, and 200 nL of AAV9 *hSyn-GRAB^DA3m^* was injected into the VS. After viral injection, the incision was sutured, and the animal was warmed using a heating pad to facilitate recovery from anesthesia. The animal was then returned to the home cage.

### Histology and histochemistry

Mice were given an overdose of isoflurane and perfused transcardially with PBS followed by 4% paraformaldehyde (PFA) in PBS. Brain tissues were post-fixed with 4% PFA in PBS overnight at 4 °C, cryoprotected with 30% sucrose solution in PBS at 4 °C for 24–48 h, and embedded in O.C.T. compound (Tissue-Tek, cat#4583). For immunostaining of GFP (Fig. S6), GCaMP6s (Figs. 1, 2, 4, and S4), YFP (Fig. 4), and Myc-tag (Fig. S4), we collected 40-μm coronal sections of the brain using a cryostat (model #CM1860; Leica). Free-floating slices were then incubated in the following solutions with gentle agitation at room temperature: 2 h in blocking solution (5% heat-inactivated goat serum, 0.4% Triton-X100 in PBS); overnight at room temperature in primary antibody 1:1000 mouse anti-GFP (GFP-1010, Aves Labs), or anti-Myc (Santa Cruz, Cat #sc-40, RRID: AB_627268) in blocking solution; 2–3 h in secondary antibody 1:500 anti-chicken-IgY Alexa488-conjugated, or goat anti-rat-IgG Cy3-conjugated or goat (Jackson ImmunoResearch) in blocking solution; 15 min in 2.5 μg/mL of DAPI (Santa Cruz, Cat #sc-3598) in PBS. Sections were mounted on slides and cover-slipped with mounting media (Fluoro-gold). Expression of stGtACR2-FusionRed (Fig. 1 and 4), hM4Di-mCherry (Fig. 7), or tdTomato (Fig. S6C) was detected through epifluorescence using the Cy3 filter without immunostaining. The expression of GRAB^DA3m^ (Fig. 6) was detected through epifluorescence using the GFP filter without immunostaining.

Sections were imaged using an Olympus BX53 microscope with a 4× (NA 0.16) or 10× (NA 0.4) objective lens equipped with a cooled CCD camera (DP80; Olympus) or Zeiss Axio Scan.Z1 with a 10× (NA 0.45) objective lens. Every third, a total of five coronal brain sections were analyzed for quantification (Fig. 1C).

### Fluorescent *in situ* hybridization (ISH)

Fluorescent ISH was performed as previously described (*77*). In brief, mice were anesthetized with isoflurane and perfused with PBS followed by 4% PFA in PBS. The brain was post-fixed with 4% PFA overnight. Thirty-micron coronal brain sections were made using a cryostat (Leica) and placed on MAS-coated glass slides (Matsunami). To generate cRNA probes, DNA templates were amplified by PCR from the C57BL/6j mouse genome or whole-brain cDNA (Genostaff, cat#MD-01). A T3 RNA polymerase recognition site (5’-AATTAACCCTCACTAAAGGG) was added to the 3’ end of the reverse primers. The primer sets to generate DNA templates for cRNA probes were as follows (the first one, forward primer, the second one, reverse primer):

### *DAT* 5’-TGCTGGTCATTGTTCTGCTC; 5’-ATGGAGGATGTGGCAATGAT

DNA templates (500–1000 ng) amplified by PCR were subjected to *in vitro* transcription with DIG (cat#11277073910)-RNA labeling mix and T3 RNA polymerase (cat#11031163001) according to the manufacturer’s instructions (Roche Applied Science).

For ISH combined with anti-GFP staining, after hybridization and washing, sections were incubated with horseradish peroxidase-conjugated anti-DIG (Roche Applied Science cat#11207733910, 1:500) and anti-GFP (Aves Labs cat#GFP-1020, 1:500) antibodies overnight. Signals were amplified by TSA-plus Cyanine 3 (AKOYA Bioscience, NEL744001KT, 1:70 in 1× plus amplification diluent) for 25 min, followed by washing, and then GFP-positive cells were visualized by anti-chicken Alexa Fluor 488 (Jackson Immuno Research cat#703-545-155, 1:250).

### Pup retrieval assay

For the ablation experiment in Figure 1A–G, animals were placed in their home cage (191 × 376 × 163 mm) with standard wood chip bedding more than a day before the retrieval assay on experimental day 0. The retrieval assay was initiated by introducing two pups (Postnatal days 1–3 at experimental day 0) in opposite corners of the nest. If the two pups were successfully retrieved, another two pups were placed in the same corners. This process continued until a total of six pups were collected or a 4-min time-out occurred. Following the retrieval assay on experimental day 0, the tested virgin female was transferred to the cage of a mother and pups to initiate cohousing. On experimental days 1 and 2, the cohoused mother and pups were temporally removed, and the assay began after a 5–10-min wait. The same mother and pups were consistently used for both the assay and cohousing across different experimental days. For example, pups of postnatal day 3 were used for the experimental day 2, if pups of postnatal day 1 were used for the experimental day 0.

For the optogenetic experiment in Fig. 1H–R, animals were placed in their small-sized home cage (191 × 188 × 163 mm) with standard wood chip bedding more than a day before the retrieval assay on experimental day 0. To habituate animals to the experimental setup, they were tethered with a patch cable (NA = 0.5, core diameter = 200 μm, TH200FL1A, Thorlabs) for 5–10 min. This habituation process was repeated more than twice before the experimental day 0. On the experimental day, the patch cable was connected to the animal, followed by a 5–10-min wait before laser illumination (IOS-465, RWD), adjusted to 5 mW/mm^2^ at each tip of the optic fiber. Immediately after the laser illumination, three pups were introduced at the three different corners, avoiding the corner where the nest had been built. After 15 min of pup exposure, a pup was removed and two pups were left in the nest. Subsequently, a retrieval assay was performed following the same procedure as described above. The same pups (postnatal day 1–3 at experimental day 0) were used across different experimental days.

For microendoscopic or photometry recordings (Figs. 2, 4, 6, 7, S2, S4, S5, and S7), animals were placed in their home cage (191 × 376 × 163 mm) with standard wood chip bedding more than a day before the retrieval assay, which was conducted once a day for 3 consecutive days. The retrieval assay was carried out in the same manner as described above, with the assay duration extended to 6 min, the length of an imaging session. We waited more than 1 min before proceeding to the next imaging session in the case of Inscopix recordings. Following the retrieval assay on the first day, the tested virgin female was transferred to the cage of a mother and pups. For subsequent days, the cohoused mother and pups were temporally removed, and after connecting the animals to the imaging rigs, a 5–10-min wait was inserted before the imaging session. The same mother and pups were used for the assay and cohousing across different experimental days. The habituation process, involving connecting animals to the imaging rigs for 5–10 min, was conducted at least twice before the first imaging. An animal showing more than five retrievals per session for the first time was defined as AP1 and the day following AP1 was defined as AP2. On the day for collecting Mother stage data, the pups were removed, leaving two to three pups in the nest. After a 2–3-min wait, an imaging session was initiated. For photometry recording with optogenetic silencing of the OFC (Figs. 4, 5, and S6), all the procedures were the same, but the duration per session was reduced to 2.5 min, and the interval between sessions was 2 min.

For microendoscopic recordings reported in Fig. 3, animals were placed in their home cage (191 × 376 × 163 mm) with standard wood chip bedding more than 1 day before the cohousing. After 2 or 3 days of cohousing, the habituation process and retrieval assay were carried out in the same manner as described above. The duration per session was 6 min, and two imaging sessions were conducted per animal.

### *In vivo* microendoscopic imaging

For microendoscopic recording, a ProView GRIN lens (500-μm diameter, 4 mm length, Inscopix) insertion was performed (OFC, coordinates relative to the bregma: AP 2.5 mm, lateral 1.2 mm, depth 1.8 mm from the brain surface). Mice were anesthetized with 65 mg/kg ketamine (Daiichi-Sankyo) and 13 mg/kg xylazine (Sigma-Aldrich) via intraperitoneal injection and head-fixed to the stereotaxic equipment (Narishige). Next, we performed a craniotomy (1 mm diameter round shape) over the lens target area, clearing any remaining bone and overlying dura using fine forceps. We aspirated the brain tissue in 1 mm. Then, a GRIN lens was loaded onto the ProView lens holder and attached to Inscopix nVista. This unit was slowly lowered into the brain while we monitored the expression of GCaMP6s through nVista. Once the intended depth was reached and the signals from GCaMP6s were confirmed, we finalized the lens placement by permanently gluing the lens with Super-Bond (Sun Medical) and sealing the lens and skull together. In this step, we also glued a metal bar (Narishige, CF-10), which allowed us to attach and detach the microscope easily. After the glue was completely hardened, the camera and lens holder were carefully released from the lens, and Kwik-Kast (WPI) was used to protect the exposed lens surface. After more than 3 weeks of recovery, the mice were anesthetized and placed in the stereotaxic equipment again. The focal plane was adjusted until GCaMP6s-labeled cells were in focus, and then a baseplate (Inscopix) was permanently glued with Super-Bond. After more than a week of recovery, we attached the microscope and let the mice explore freely in their home cage for 5–10 min. We performed this habituation session more than twice before the first imaging session.

We performed microendoscopic imaging using the Inscopix nVista system (Inscopix). We performed imaging without refocusing the microscopes across imaging sessions during the day, whereas the focal plane was adjusted each day before the first imaging session. Before imaging, we attached the microendoscope to the animals while holding the implanted head bar. Images (1080 × 1080 pixels) were acquired using nVista HD software (Inscopix) at 10 Hz, with LED power of 0.4– 1 mW/mm^2^ and a gain of 2.0–3.0. Time stamps of the imaging frames, camera, delivery of sucrose water were collected for alignment using WaveSurfer (https://wavesurfer.janelia.org/). We performed two imaging sessions, each lasting for 6 min, with an interval of a few minutes between sessions. The imaging data were cropped to 800 × 700 pixels and exported as .tiff files using the Inscopix Data Processing Software. To identify ROIs corresponding to putative cell bodies for the extraction of neural signals, we used v2 of MIN1PIPE (https://github.com/JinghaoLu/MIN1PIPE (*78*)) with a spatial down-sampling rate of 2. All traces from identified ROIs were manually inspected to ensure quality signals and excluded if they had an abnormal shape or overlapped signal from adjacent ROIs. Relative changes in calcium fluorescence *F* were calculated by *dF/F_0_ = (F-F_0_)/F_0_* (where *F_0_* is the median fluorescence of the entire trace). *dF/F* was normalized within each cell. In the field of view, we typically detected 50.8 ± 11.1 (mean ± SD) ROIs.

Behavior videos were acquired at 20 Hz using a camera (Imaging source, DMK 33UX174). WaveSurfer was used to generate precise transistor-transistor logic (TTL) pulses to synchronize behavioral tracking and microendoscopic imaging. The animals that showed 10 or more successful pup retrievals were included in the following analysis.

For longitudinal tracking of the same cells across different days (Figs. S3 and S5), spatial footprints extracted using the MIN1PIPE pipeline were processed with CellReg (*79*) using default parameters (maximum translation: 12 microns, registration threshold P_same = 0.5). The AP2 session was used as the reference for filed-of-view alignment.

### Clustering analysis

We performed clustering analysis on the averaged responses during pup retrieval trials to identify functional subtypes (clusters) and evaluate heterogeneity in OFC^Rbp4^ neuron activity. For this, activity traces from –2 to 8 seconds relative to pup contact were averaged per trial. Data were pooled across AP1, AP2, and Mother stages, resulting in a 1232 × 100 data matrix. Principal component analysis (PCA) was applied using MATLAB’s *pca* function to reduce dimensionality. The first six principal components (PCs), which together explained over 95% of the variance (Fig. S2), were selected. *K*-means clustering was then performed on the PC scores using six clusters and a fixed random seed for reproducibility. To classify new samples into existing clusters, we normalized the new data using the mean of the original dataset to ensure centering consistency. The centered data were projected onto the original PCA space using the original PC vectors. Cluster assignment was then performed using the *knnsearch* function in MATLAB to find the nearest cluster centroids.

To quantify response magnitude (Figs. 2, 5, S4, S5, and S7), we used the following time windows relative to pup contact: Cluster 1: –1 to +1 s; Clusters 2, 3, and 5: 0 to +4 s; Cluster 4: 0 to +6 s. Baseline windows were as follow: Cluster 1: –3 to –2 s; Clusters 2, 3, 4, and 5: –2 to –0.5 s. Response amplitudes were quantified as the AUC during the response window, relative to the mean baseline activity for each cluster.

### Decoder analysis

To decode behavioral events among three categories (pup contact, onset of retrieval, and completion), we trained a set of binary SVM classifiers using a one-vs.-one multiclass strategy implemented via MATLAB’s *fitcecoc* function (Fig. 2J, K). Data from the initial 10 retrieval trials were used for both training and testing, with the number of trials adjusted to match the minimum across conditions. For each iteration, the dataset was split into training and test sets at a 4:1 ratio. Calcium activity was convolved over a 500-ms time window from the onset of each behavioral event. The training involved fivefold cross-validation, and classifier outputs were aggregated by majority voting to determine the most likely behavioral label. Decoding accuracy was quantified as the percentage of correctly classified test samples. Each classification run was repeated 5,000 times, with the mean decoding accuracy computed across iterations. In each run, different subset of ROIs were randomly selected for training.

To decode whether a pup contact led to a retrieval or non-retrieval outcome (Fig. 2L), we trained a binary SVM classifier using the MATLAB’s *fitcsvm* function. The classifier was trained and tested using data from the first six retrieval and six non-retrieval trials. In each run, data were split into training and test sets at a 2:1 ratio. Threefold cross-validation was applied during training, and Ca^2+^ signals were convolved over a 500-ms window from the moment of pup contact. Classification accuracy was calculated as the percentage of correctly identified trials in the test set. This process was repeated 10,000 times, with the mean accuracy determined across iterations. Subsets of cells were randomly selected in each run.

### PCA trajectory analysis

To assess the ability of neural population activity to discriminate between retrieval and non-retrieval trials, we performed PCA on simultaneously recorded ROIs from the AP1, AP2, and Mother stages, and calculated pairwise Euclidean distances. The analysis was conducted using MATLAB’s *pca* function. For each population of simultaneously recorded ROIs, a time series was constructed by concatenating all retrieval and non-retrieval trials into an *n* × (*n*-trials × time-bins-per-trial) matrix, where *n* represents the number of ROIs. The time-bins-per-trial was set to 80, corresponding to a 100-ms bin width across an 8-second window centered on pup contact (–2 to +6 seconds). PCA was applied to this concatenated dataset, and the first three principal components (PCs) were extracted. Each trial’s time series was projected onto these three PCs to generate a trajectory in PCA space. Mean trajectories were then computed across trials for each behavioral condition (Fig. 2N).

### Nonsocial reward presentation

10% sucrose water was delivered from a behavioral lick port (Sanworks) controlled by a MATLAB-based open-source state machine (Bpod; Sanworks). Mice could access the sucrose water whenever they broke an IR beam in front of the lick port and waited for longer than 1 s. To habituate the mice to the setup, the lick port was introduced one day before the imaging session. After a habituation period exceeding 12 hours, mice were water-deprived for 12–16 hours to enhance the value of water. Then, we performed imaging during a pup retrieval assay, in which mice could freely access to the licking port. Each animal underwent two to four imaging sessions (duration 6 min). TTL pulses from the state machine were used to synchronize the timing of water delivery and microendoscopic imaging. To define significantly water-responsive ROIs, trial-averaged Ca^2+^ signals were compared between the licking event and a pre-event baseline using the Mann–Whitney *U* test, with a significance threshold of p < 0.05.

### Test for significant overlap between nonsocial reward and pup-retrieval responses

To evaluate the significance of the overlap between ROIs responsive to water reward and pup retrieval, we compared the observed proportion of dual-responsive ROIs with the overlap expected by chance. The chance overlap level was estimated using a null distribution created through the following procedure (*80*): the labels indicating responsiveness to water reward and pup retrieval were randomly shuffled across all imaged ROIs, while maintaining the total number of ROIs responsive to each event. For each permutation, the percentage of ROIs labeled as responsive to both events was calculated. This randomization was repeated 5,000 times to construct a distribution of expected overlap under the null hypothesis. The significance of the observed overlap was determined by comparing it to this simulated distribution.

### Fiber photometry with optogenetic stimulation

For fiber photometry recording (Figs. 4, 5), *DAT-Cre/+; Ai162/+* double heterozygous female mice were used. We implanted the optical fibers (NA = 0.50, core diameter = 200 μm, 3 mm length from RWD) into the bilateral OFC for optogenetic stimulation at 2 weeks after AAV injection into the OFC (coordinates relative to the bregma: anterior 2.5 mm, lateral 0.6 mm, depth 1.7 mm from the brain surface with tilting 5° from the vertical). At the same surgery, an optical fiber (NA = 0.50, core diameter = 400 μm from Kyocera) was implanted above the VTA for fiber photometry. After surgery, the animals were crossed with stud males and housed in the home cage until recording. We performed Ca^2+^ imaging by delivering excitation lights (470 nm modulated at 530.481 Hz and 405 nm modulated at 208.616 Hz) and collected the emitted fluorescence using the integrated Fluorescence Mini Cube (Doric, iFMC4_AE(405)_E(460-490)_F(500-550)_S). Light collection, filtering, and demodulation were performed using the Doric photometry setup and Doric Neuroscience Studio Software (Doric Lenses, Inc.). The 405-nm signal was recorded as a background (non-calcium-dependent), and the 470-nm signal reported calcium-dependent GCaMP6s excitation/emission. The power output at the tip of the fiber was about 50 μW. The signals were initially acquired at 12 kHz and then decimated to 120 Hz. Next, we down-sampled the raw Ca^2+^ data to 20 Hz using the MATLAB’s *resample* function.

For optogenetic stimulation, animals were connected to a 465-nm laser (RWD) via split optical patch cords (NA = 0.50, core diameter = 200 μm, 200TH200FL1A; Thorlabs) and a rotary joint (RJ1; Thorlabs). For optogenetic inhibition of the OFC using GtACR2, 150 s of 465 nm continuous photostimulation at 5 mW/mm^2^ at the fiber tip was used during the pup retrieval session. Of note, previous study showed that 30 minutes of continuous light stimulation in GtACR2-expressing VTA^DA^ neurons sufficiently induce place avoidance (*81*). We first performed a photometry imaging session without light stimulation, followed by a session with light stimulation, which was repeated twice.

### Fiber photometry with a GRAB^DA^ sensor

For fiber photometry recording of DA dynamics (Fig. 6 and 7), a custom-built photometry system was used. The light from the fiber-coupled light-emitting diode (LED; 470 nm, M470F4, Thorlabs) was collimated (F950FC-A, Thorlabs) to pass through an excitation filter (MDF-GFP2 482-18, Thorlabs) and dichroic mirrors (MDF-GFP2, Thorlabs; ZT/405/488/561/647, Chroma). The filtered light was focused onto a fiber-optic patch cable (NA = 0.50, core diameter = 400 μm, MAF2L1, Thorlabs) through a rotary joint (RJ1; Thorlabs). The power output at the tip of the fiber was adjusted to 5 to 10 μW. The patch cable was connected to the optic fiber (NA = 0.50, core diameter = 400 μm, 6 mm length from RWD) implanted in the VS, DS, pDLS, or BLA. For isosbestic excitation at 405 nm, another light from the fiber-coupled LED (405 nm, M405FP1, Thorlabs) was collimated (F950FC-A, Thorlabs) and passed through an excitation filter (MF390-18, Thorlabs) and a dichroic mirror (MD416, Thorlabs) to merge with the 470 nm light path. The emission fluorescence was detected using a photomultiplier tube (PMT1001/M, Thorlabs) after passing through the dichroic mirror (ZT/405/488/561/647, Chroma) and an emission filter (MDF-GFP2 520-28, Thorlabs). LEDs were alternately illuminated for 4 ms with 6-ms intervals triggered by an open-source pulse generator (Pulse Pal v2, Sanworks) to detect signals at 470 nm and 405 nm separately. PMT signals were recorded at 1 kHz using a data acquisition system (PCIe-6341, National Instruments) and synchronized with a camera through WaveSurfer. The mean value of the middle 2-ms period during 4-ms LED illumination was used for analysis, which resulted in a 100 Hz signal. Signals were further decimated to 20 Hz after individual channel smoothing using the MATLAB *smooth* function with a LOWESS local linear regression method.

### Photometry data analysis

We analyzed the photometry data using custom-written MATLAB codes. To calculate dF/F, a least-squares linear fit was applied to the 405-nm signal to align it to the 470-nm signal, producing a fitted 405-nm signal that was used to normalize the 470-nm signal using the MATLAB *polyfit* function. The dF/F was generated by subtracting the fitted 405-nm signal from the 470-nm signal to eliminate movement or other common artifacts. Finally, Ca^2+^ or DA traces were Z-scored by the mean and standard error of the traces for an entire recording session. To calculate the averaged response, we took a 2.5-s time window (0–2.5 s for Ca^2+^ and –0.5–2 s for DA traces following pup contact), and that value was subtracted by the mean of –2 to –0.5 s preceding the pup contact as a baseline.

To determine the specificity of DA transients in response to pup retrieval (Fig. 6E), we measured the AUC-ROC. This was calculated by comparing the distribution of Ca^2+^ responses for each time frame along the trial (mean response for 10 frames, equivalent to 0.5 s from the time point) versus the distribution of calcium responses for the baseline (mean response from –3 to – 2 s before pup contact). The AUC-ROC value ranges from 0 to 1 and quantifies the accuracy of an ideal observer. Values proximal to 0.5 indicate low discrimination, whereas values far from 0.5 indicate high discrimination relative to baseline activity. To assess significance, we calculated the sample distribution by temporal shuffling within trials (n = 10 iterations). The timings where the signal exceeded the mean + 2SD of the sample distribution were defined as significant (Fig. 6E, red dots).

### Pharmacogenetics

To activate (Fig. S4) or silence (Fig. 7) neural activity by hM3Dq or hM4Di, respectively, 100 μL of CNO (0.5 mg/mL, Sigma; C0832) was intraperitoneally injected after two 6-min imaging sessions (pre-CNO sessions). The two imaging sessions were performed 30 min after the injection of CNO (post-CNO sessions).

### Counting the number of GtACR2+ cells

To quantify the number of GtACR2+ cells (Fig. 1N), we utilized a semi-automated approach using ilastik (*82*) in combination with custom MATLAB code. We trained ilastik on several slices to detect GtACR2+ cells. The trained classifier was then used to generate binary masks for the GtACR2 channel. To reduce noise, we applied MATLAB’s morphological opening function to the binary masks. The number of GtACR2+ cells was counted bilaterally beneath the fiber tracts. For each mouse, three slices containing the fiber tract were analyzed.

### Quantification of the overlap between OFC^Rbp4^ neurons and VTA-projecting neurons

To evaluate the overlap between VTA-projecting neurons in the OFC and OFC^Rbp4^ population (Fig. S6C–E), we manually counted tdTomato+ and tdTomato+ GCaMP6+ dual-labeled cells using custom MATLAB code. For each mouse, five slices were analyzed (every other section), ensuring consistent sampling across animals.

### Analysis of locomotor activity

To quantify locomotor activity (Figs. S6H and S8), we tracked mouse body trajectories using SLEAP (*83*). A SLEAP model was trained to identify the nose, head, left and right ears, and back, based on manually labeled frames selected randomly from the analyzed videos. The mean coordinates of all detected body parts were used to represent the position of the mouse in each frame, accounting for variability in part detection. Locomotor activity was quantified as the frame-to-frame displacement of the calculated body position.

### Quantification and statistics

Statistical tests were performed using custom-written MATLAB codes. All tests were two-tailed. The sample size and statistical tests used are indicated in the figures or corresponding legends. The criterion for statistical significance was set at P < 0.05. The mean ± SEM was used to report statistics unless otherwise indicated. All of the tests that were used in this study and their p-values are summarized in the Supplementary Table (Table S1).

## Acknowledgments

We thank Liqun Luo, Adi Mizrahi, Ido Maor, and members of the Miyamichi lab for the critical reading of the manuscript, the University of North Carolina vector core and the Canadian Neurophotonics Platform viral vector core facility for the AAV productions, Masanori Murayama for sharing the *Rbp4-Cre* mice, Noriaki Ohkawa for the advice on the miniaturized microscope, Kentaro Ishii for the advice on the decoder analysis, and the animal facility of RIKEN BDR for taking care of the animals and *in vitro* fertilization.

## Funding

JST PRESTO program JPMJPR21S7 (GT)

JSPS KAKENHI 20K15941 (GT)

RIKEN Incentive grant (GT)

RIKEN Diversity grant (GT)

AMED Brain/MINDS program JP21dm0207113 (KaK)

JSPS KAKENHI 21H02587 and 23H04945 (KM)

Takeda Science Foundation Research Grant (KM)

RIKEN BDR Stage Transition Project (KM)

## Author contributions

Conceptualization: GT, KM

Methodology: GT, MH, SI, KI, KeK, SK, KaK

Investigation: GT, MH, SI, HK

Supervision: KM

Writing—original draft: GT, KM

Writing—review & editing: GT, KM

## Data and materials availability

Custom MATLAB scripts supporting the findings of this study and data from fiber photometry and microendoscopy were deposited in the SSBD repository (https://doi.org/10.24631/ssbd.repos.2024.10.402). All other data and materials are available in the main text and supplementary materials.

## Supplementary Figures

**Fig. S1.**
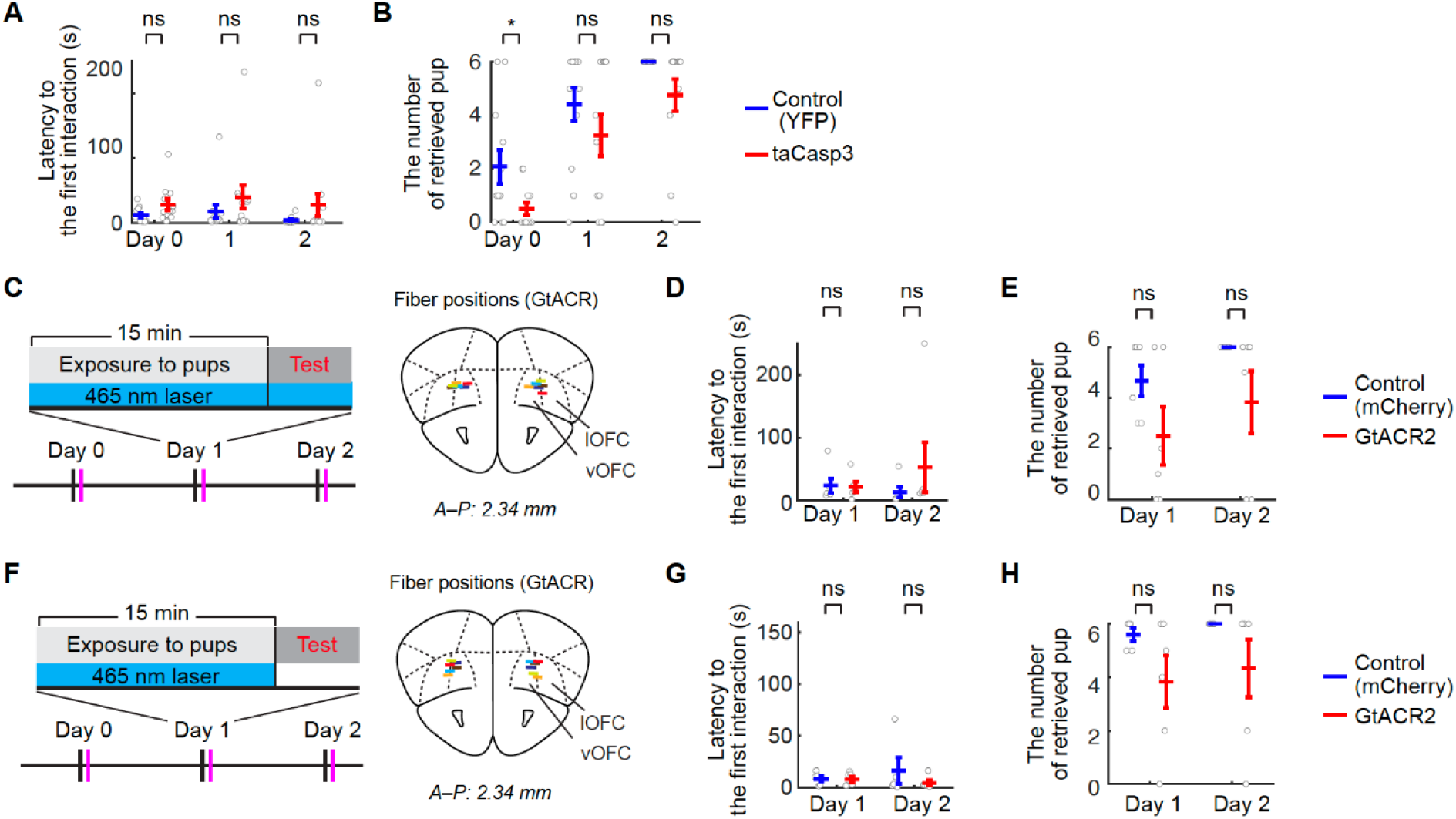
Additional data on the necessity of OFC^Rbp4^ neurons for effective acquisition of pup retrieval, related to Fig. 1. (A) Latency to the first contact. ns, not significant by two-way ANOVA with repeated measures; condition effect, ns; time course effect, ns; interaction effect, ns. (N = 12 per group). (B) Number of retrieved pups during a 4-min trial. Two-way ANOVA with repeated measures; condition effect, p < 0.01; time course effect, p < 0.001; interaction effect, ns. *p < 0.05 by post-hoc unpaired t-test with Benjamini-Hochberg correction. (N = 12 per group). (C, F) (Left) Schematic of the experimental design. (Right) Schematic coronal sections showing optic fiber placements in the GtACR2-injected group (N = 6 mice). (D, G) Latency to the first contact. ns, not significant by two-way ANOVA with repeated measures; condition effect, ns; time course effect, ns; interaction effect, ns. (N = 6 per group in panel D; N = 5 and 6 in panel G). (E, H) Number of retrieved pups in a 4-min trial. ns, not significant by post-hoc unpaired t-test with Benjamini-Hochberg correction after significant by two-way ANOVA with repeated measures; condition effect, p < 0.05; time course effect, ns; interaction effect, ns. (N = 6 per group in panel E, N = 5 and 6 in panel H).

**Fig. S2.**
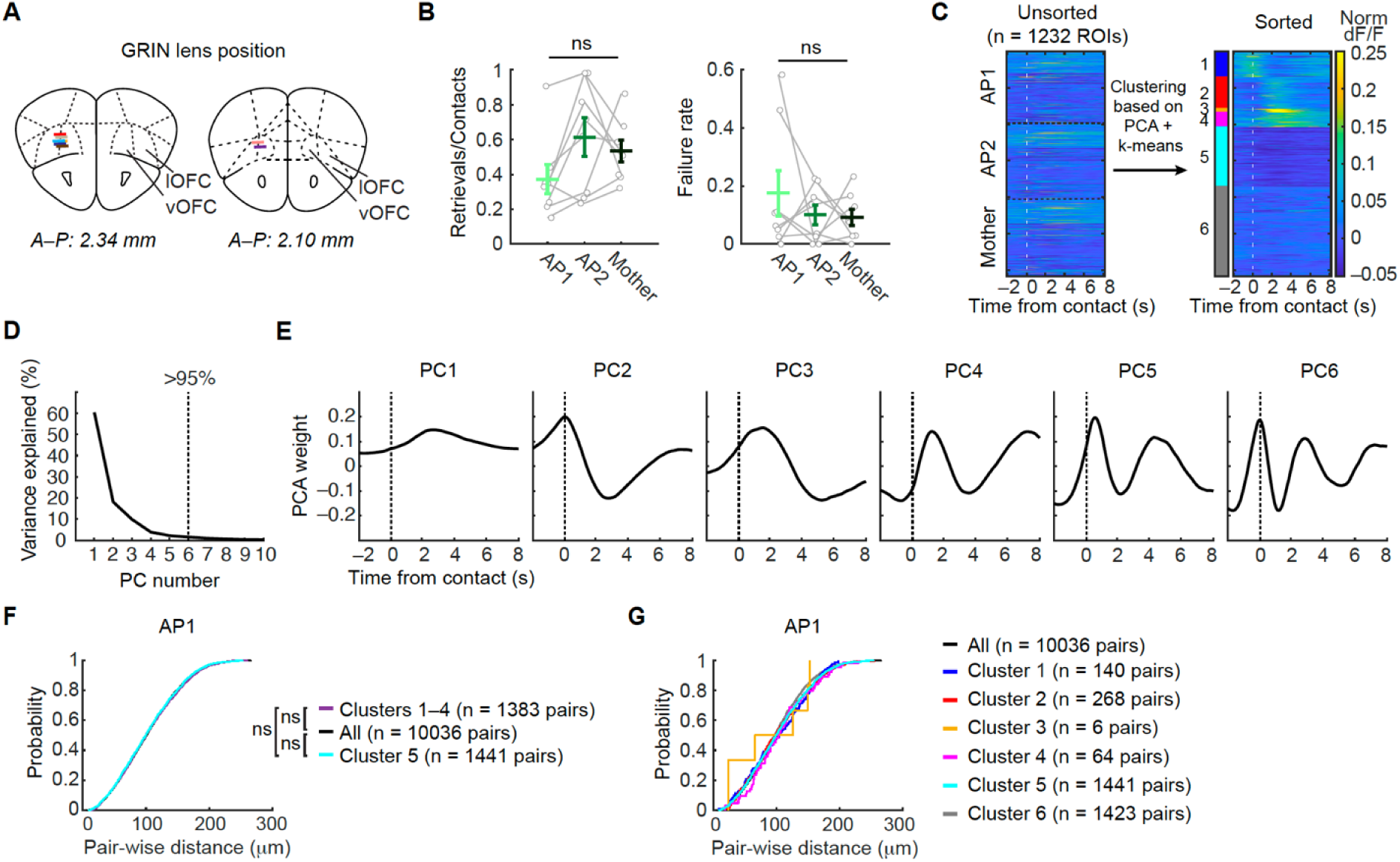
Additional data on the neural representations of OFC^Rbp4^ neurons during pup retrieval, related to Fig. 2. (A) Schematics of coronal sections showing the GRIN lens placements (N = 8 mice). (B) Quantification of pup retrieval performance during two 6-min imaging sessions (dataset includes eight mice successfully imaged across the AP1, AP2, and Mother stages). (From left to right) Success rate calculated as the number of retrievals over the number of pup contacts. Failure rate calculated as the number of incomplete retrievals (dropping before reaching the nest) normalized to the total number of retrievals. ns, not significant by a significant one-way ANOVA (N = 8 mice). Error bars indicate the SEM. (C) Normalized, averaged responses of ROIs during retrieval trials before (left) and after (right) clustering. (D) Scree plot showing the percentage of explained variance per principal component. Over 95% of the variance was accounted for by the first six principal components (dashed line). (E) Individual principal components retained for clustering, displayed as response vectors. (F) Pairwise distances among all ROIs, within clusters 1–4 (elevated responses), and cluster 5 (suppressed responses). Data are from all female mice (N = 8) on AP1. No topographical organization was found (Kolmogorov–Smirnov test; ns, not significant). (G) Pairwise distances among all ROIs and within each cluster.

**Figure S3.**
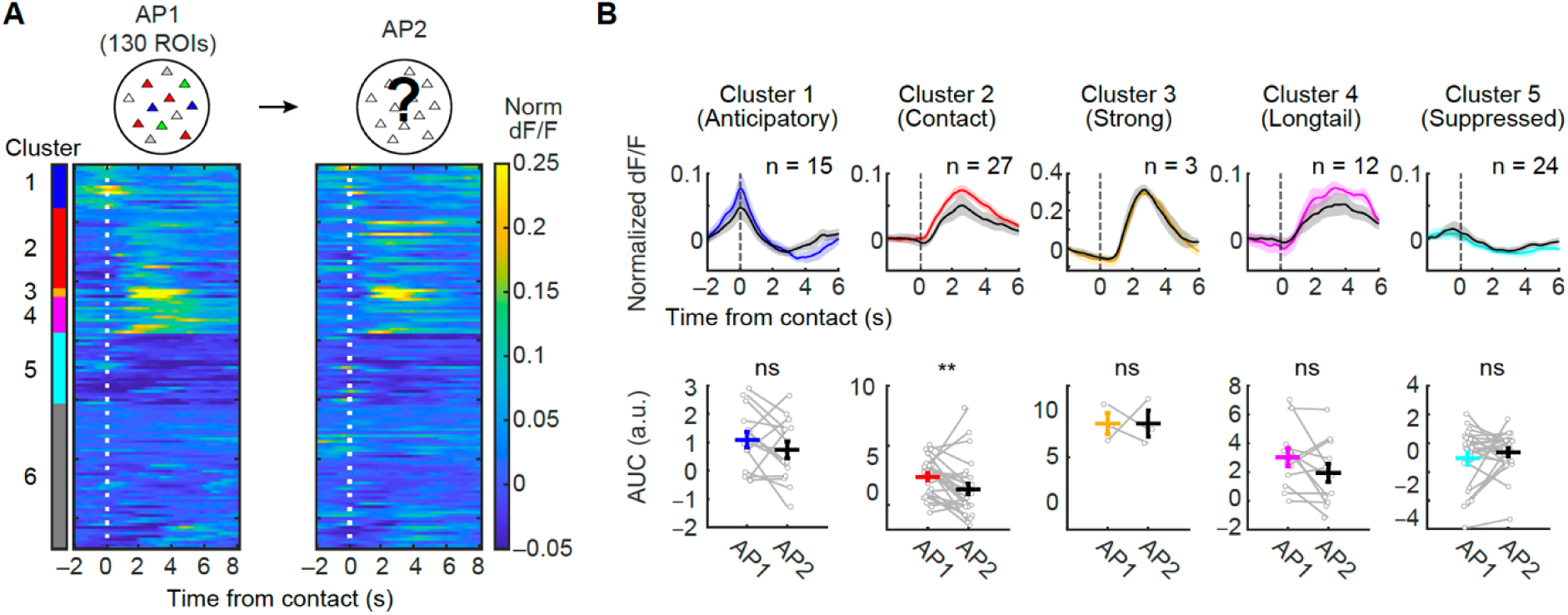
Longitudinal tracking of OFC^Rbp4^ neuron activity across days, related to Fig. 2. (A) Activity heat maps show normalized, trial-averaged longitudinal tracking of the same ROIs from AP1 to AP2. ROIs are sorted by their cluster identity on AP1. Time 0 marks pup contact followed by retrieval. (B) Quantification of mean response intensity between AP1 and AP2. **, p < 0.01 by the paired t-test. ns, not significant. The number of ROIs is indicated within the panel.

**Fig. S4.**
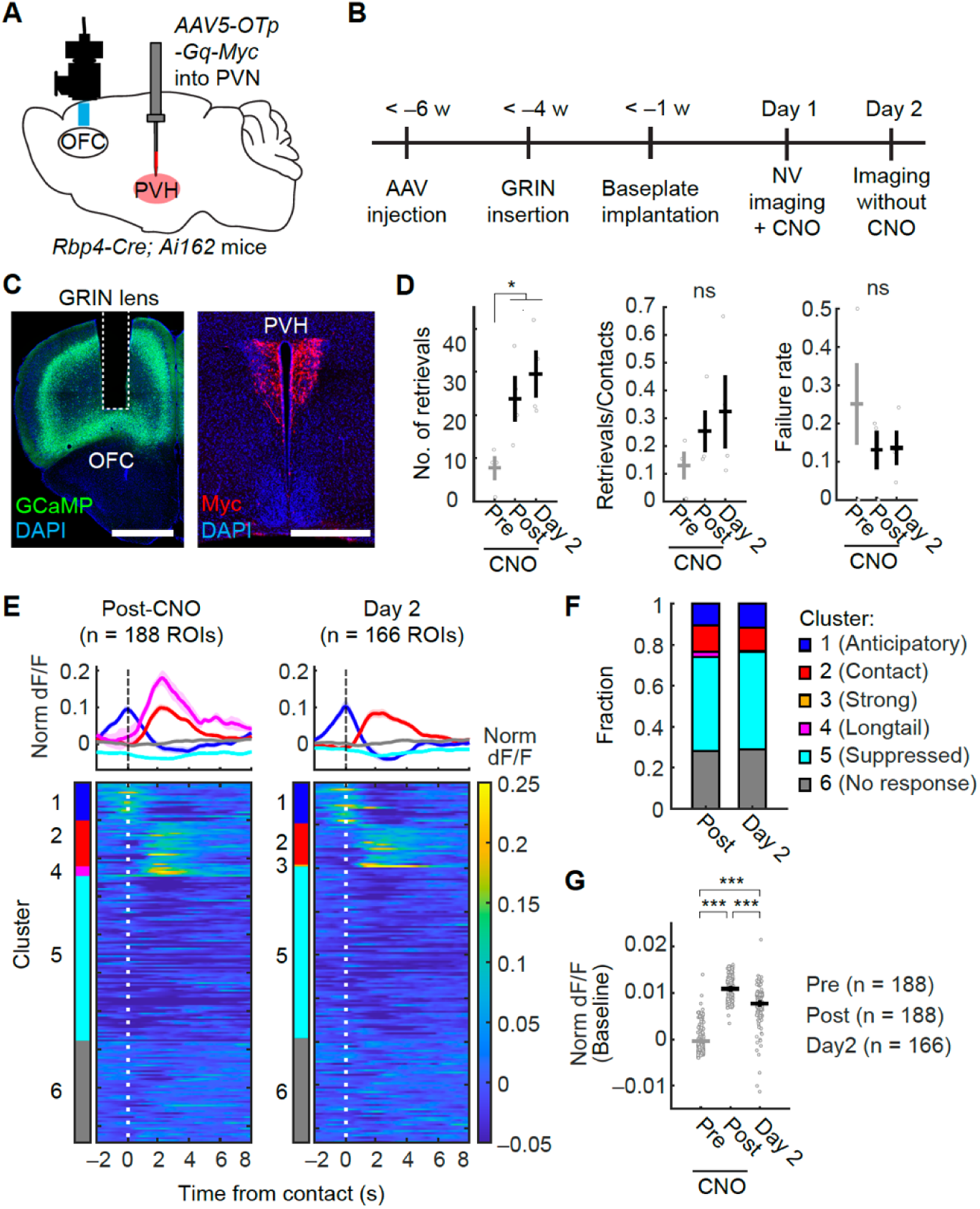
Representation of pup retrieval emerges by activating OT neurons in the PVN of virgin female mice, related to Fig. 2. (A) Schematics of the experimental design. AAV5 *OTp-hM3D(Gq)-Myc* was injected into the bilateral PVN of virgin *Rbp4-Cre; Ai162* mice. (B) Schematic of the experimental timeline. (C) Representative coronal sections showing the tract of the GRIN lens in the OFC (left) and expression of Gq-Myc in the PVN (right). Scale bars, 500 μm. (D) Quantification of pup retrieval performance during the imaging sessions. (Left) Number of pup retrievals. (Middle) Success rate as in Fig. S2B. (Right) Failure rate as in Fig. S2B. *, p < 0.05 by a post hoc Dunnett’s test after a significant one-way ANOVA (N = 4 mice). ns, not significant. Error bars indicate the SEM. (E) (Top) Trial-averaged activity traces of ROIs for each cluster during post-CNO and the subsequent Day 2 session. (Bottom) Activity heat maps showing normalized, averaged responses of individual ROIs during pup retrieval. ROIs are sorted by cluster identity, with time 0 indicating pup contact followed by retrieval (n = 188 ROIs from N = 4 mice). Cluster annotation is based on principle components and clustering space defined in Fig. S2. (F) Fraction of ROIs assigned to each cluster. (G) Quantification of baseline activity levels, calculated from a time window between –2 and –1 s relative to pup contact during non-retrieval trials. ***, p < 0.001 by a post hoc Tukey’s HSD test following a significant one-way ANOVA. The number of ROIs is indicated in the panel.

**Fig. S5.**
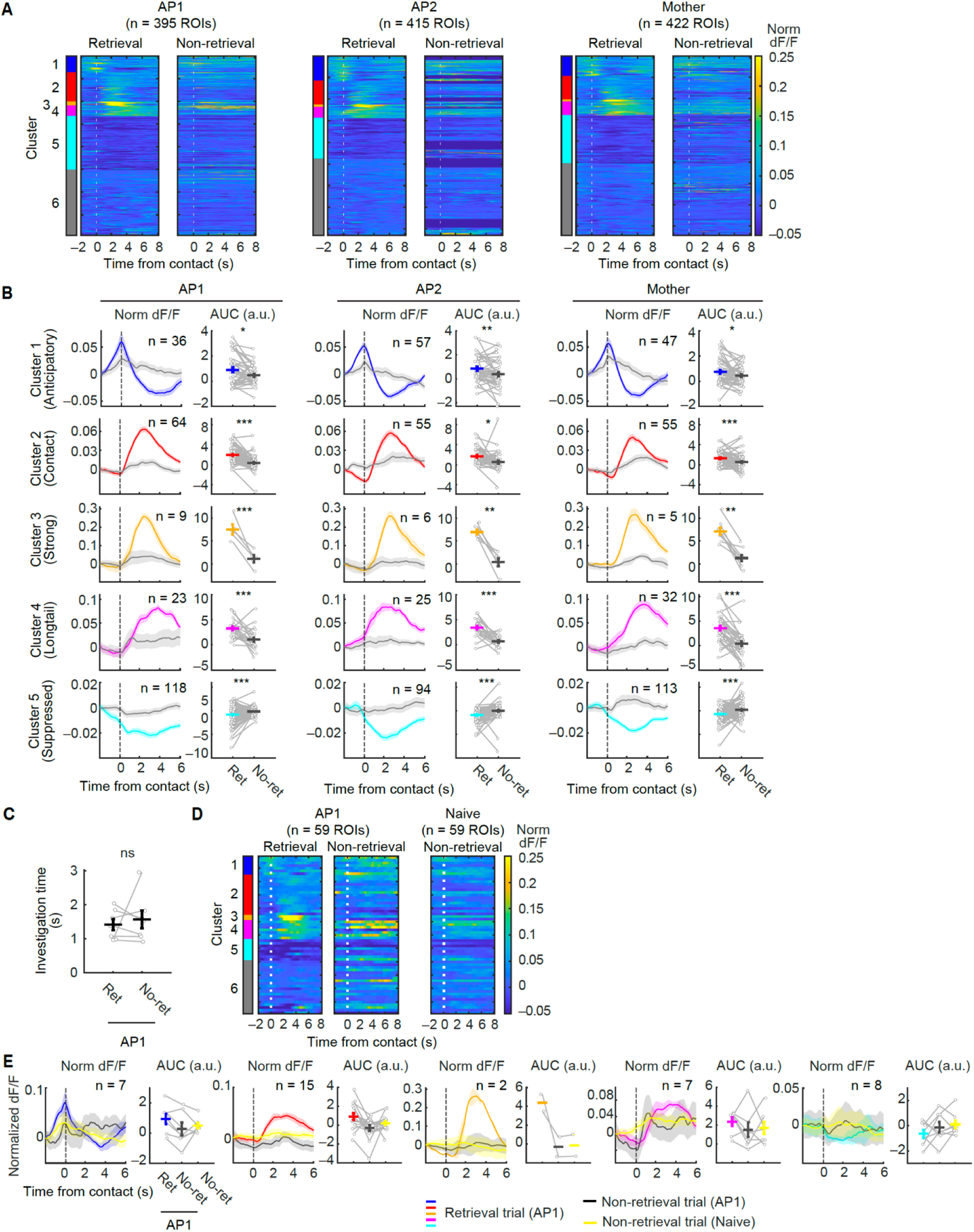
The response of OFC^Rbp4^ neurons during non-retrieval trials, related to Fig. 2. (A) Activity heat maps showing normalized, averaged responses of individual ROIs during the trials with pup retrieval (left, the same data as Fig. 2H) and contact trials that were not followed by pup retrieval (right, “Non-retrieval”). Time 0 indicates pup contact. To avoid a low statistical power, data of ROIs from the mice that performed seven or fewer non-retrieval contact trials were excluded (shown in dark blue rows). (B) Averaged activity traces of each cluster (left) and quantification of the averaged responses (right). *, p < 0.05, **, p < 0.01, ***, p < 0.001 by two-sided paired t-test. ns, not significant. (C) Quantification of pup sniffing time during retrieval and non-retrieval trials in AP1 (N = 8 mice). ns, not significant by the Wilcoxon signed-rank test. (D) Activity heat maps showing normalized, averaged responses of individual ROIs identified as the same neurons across Naïve (without experience of co-housing) and AP1 stages, during retrieval and non-retrieval trials. (E) Mean responses during retrieval trials in AP1 (colored), non-retrieval trials in AP1 (gray), and non-retrieval trials in Naïve stages (yellow). The number of datasets is indicated within the panel.

**Fig. S6.**
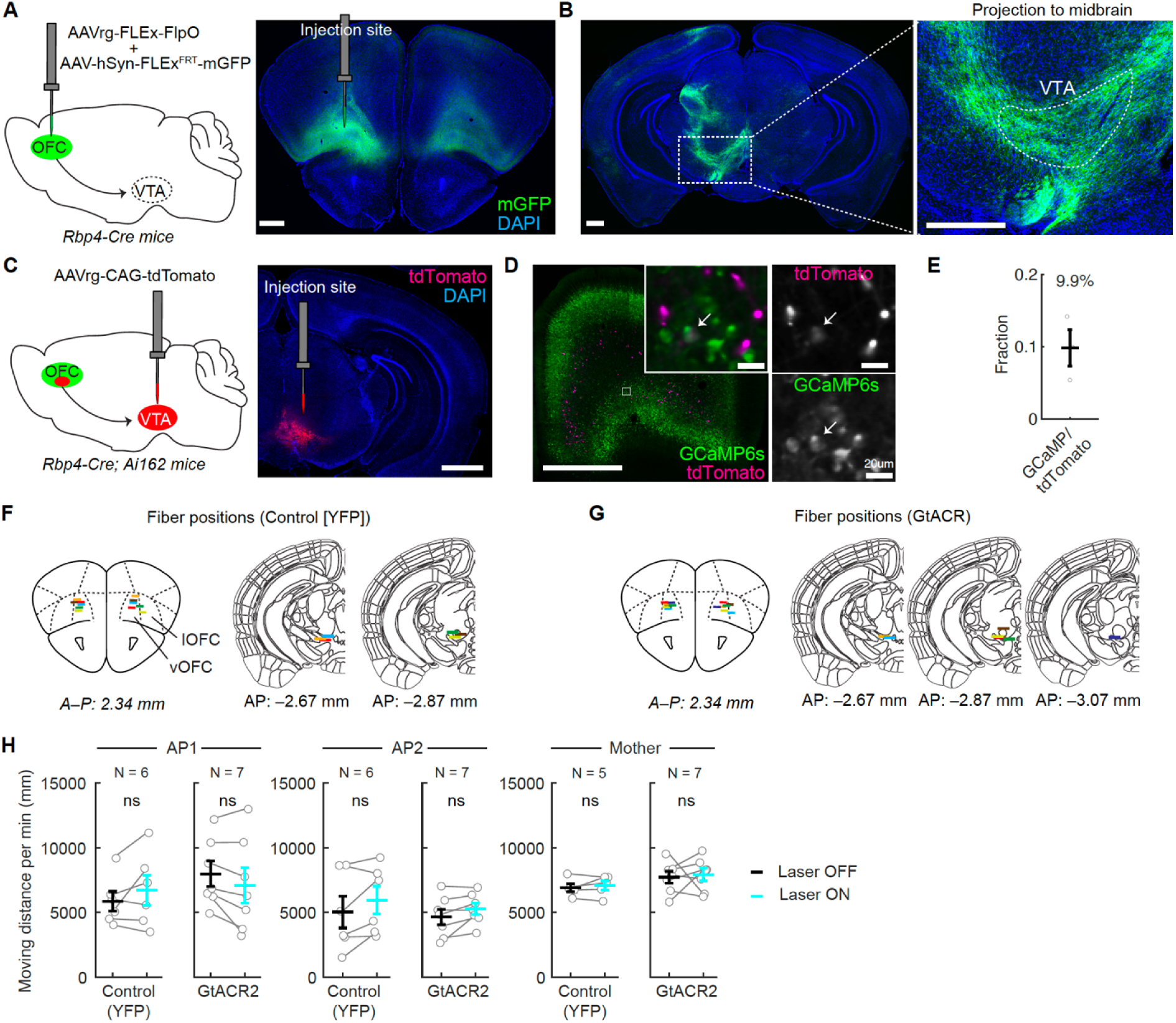
OFC^Rbp4^ neurons project axons to the VTA, and optogenetic inactivation of OFC^Rbp4^ neurons did not affect locomotor activity, related to Fig. 4. (A) (Left) Schematic of the virus injection strategy. A mixture of AAVrg *CAG-FLEx-FlpO* and AAV9 *hSyn-fDIO-mGFP* was injected into the left OFC. (Right) Representative coronal section showing the injection site. Scale bar, 500 μm. (B) Representative coronal section showing axonal projections from the OFC to the midbrain. Scale bars, 500 μm. (C) (Left) Schematic of the virus injection strategy for AAVrg *CAG-tdTomato* into the left VTA. (Right) Representative coronal section showing the injection site. Scale bar, 1 mm. (D) Representative coronal section showing the overlap between GCaMP6s (putative OFC^Rbp4+^ cells) and tdTomato (putative VTA-projecting neurons). Scale bars, 1 mm (main) and 20 μm (insets). (E) Quantification of the fraction of OFC^Rbp4^ cells that project to the VTA (N = 3 mice). (F, G) Schematic of coronal sections showing the optic fiber positions in the YFP-(N = 6 mice) and GtACR2-injected (N = 7 mice) groups. Colors indicate individual mice. (H) Quantification of locomotion activity during light off (black) and light on (cyan) trials. ns, not significant by the paired t-test. The number of animals is indicated in the panel.

**Fig. S7.**
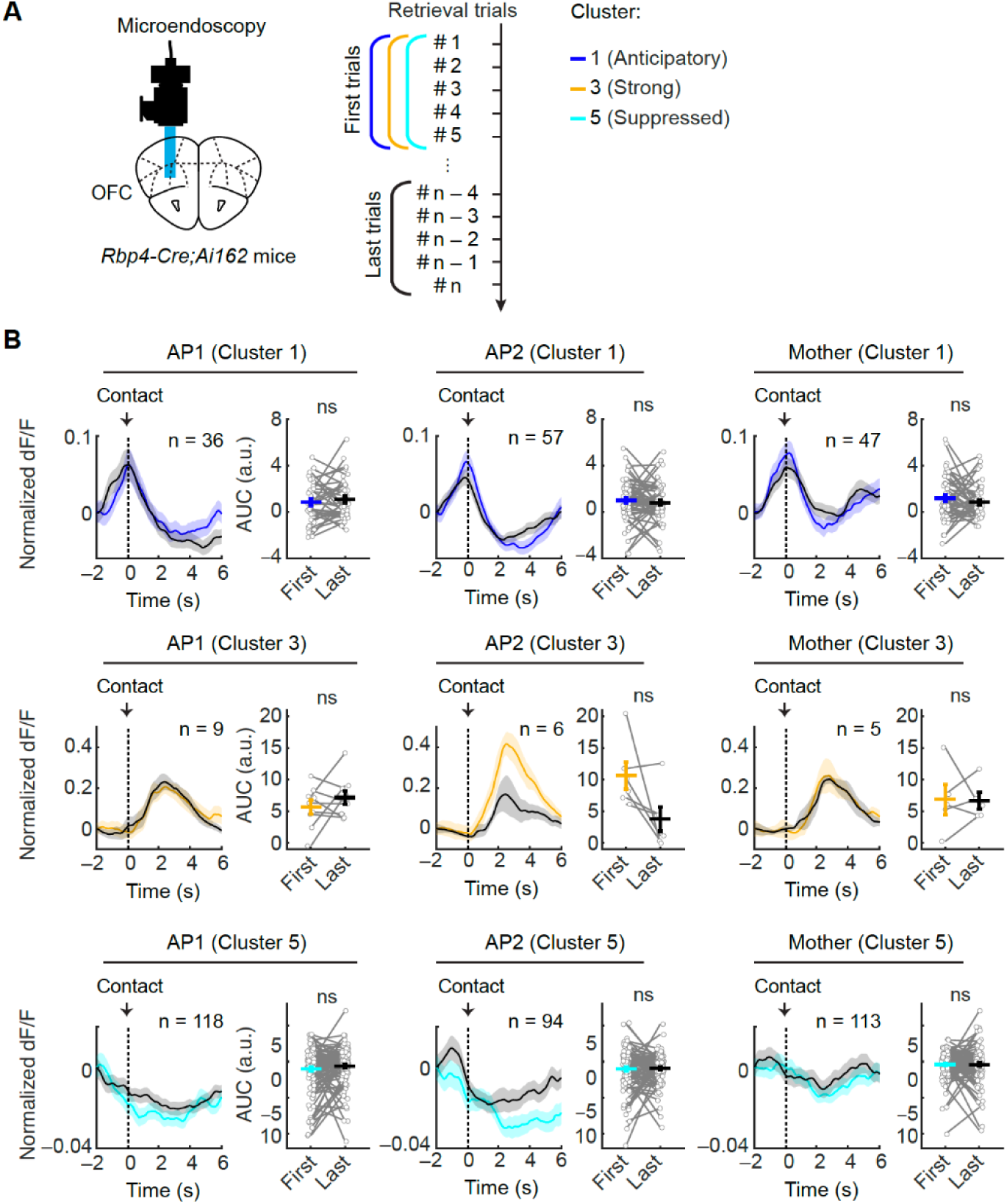
Adaptation effect in different clusters of OFC^Rbp4^ neurons, related to Fig. 5. (A) Schematic of the experimental design and analyzed trials. (B) Averaged activity traces of ROIs from each cluster for the first (colored) and last (black) five trials of pup retrieval. Time 0 indicates pup contact followed by retrieval. The shadow represents the SEM. The right graph shows the AUC of normalized dF/F between –1 and 1 s (cluster 1), and 0 and 4 s (clusters 3 and 5). *, p < 0.05 by the paired t-test. The number of ROIs is indicated in the panel.

**Fig. S8.**
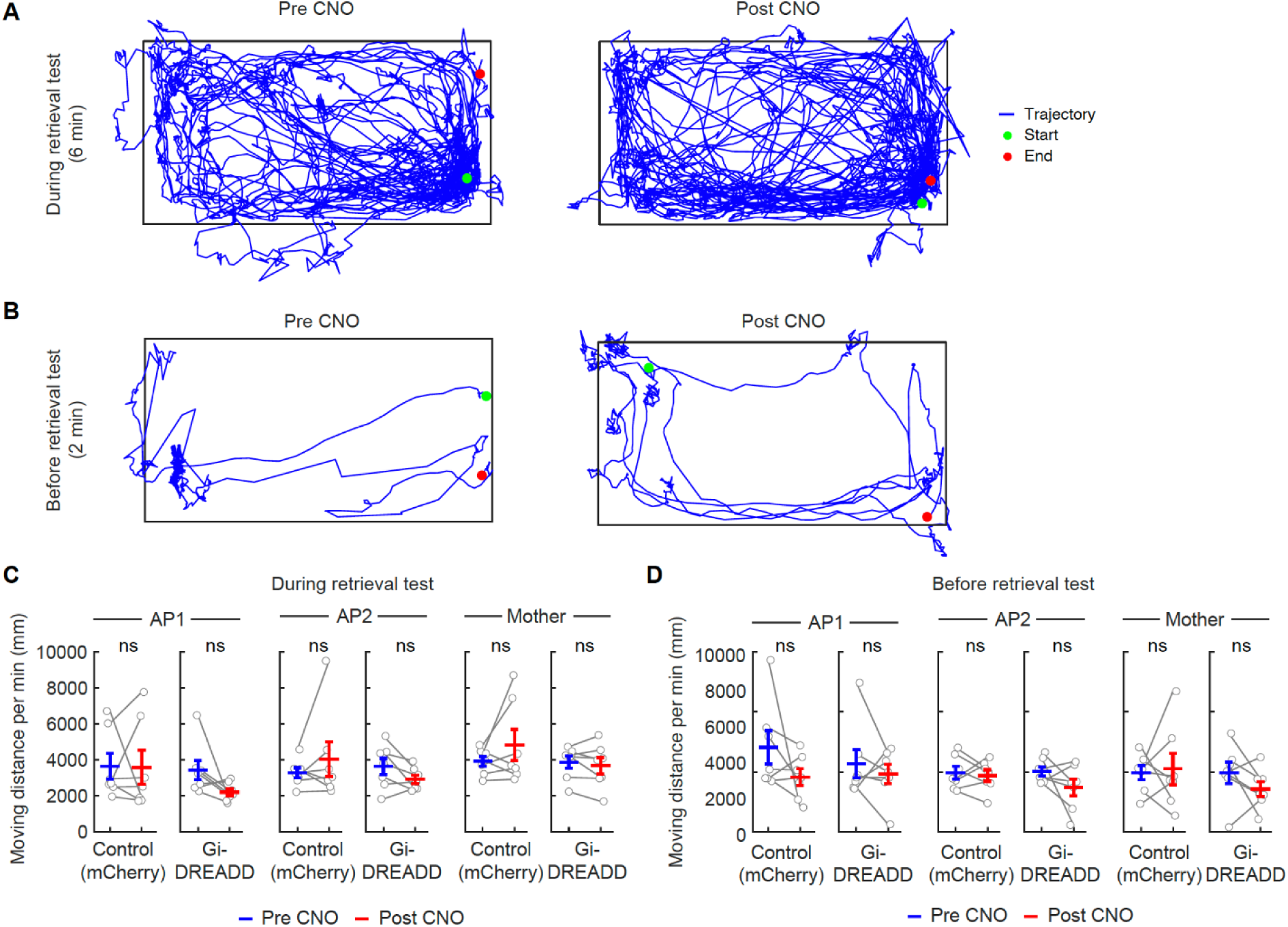
CNO treatment did not affect locomotor activity, related to Fig. 7. (A, B) Trajectories of one representative mouse during a 6-min retrieval trial before and after CNO injection (A), and baseline activity during a 2-min pre-trial period (B). (C, D) Quantification of locomotor activity during the retrieval trial (C) and pre-trial (D). ns, not significant by paired t-test (N = 7 mice per group).

**Fig. S9.**
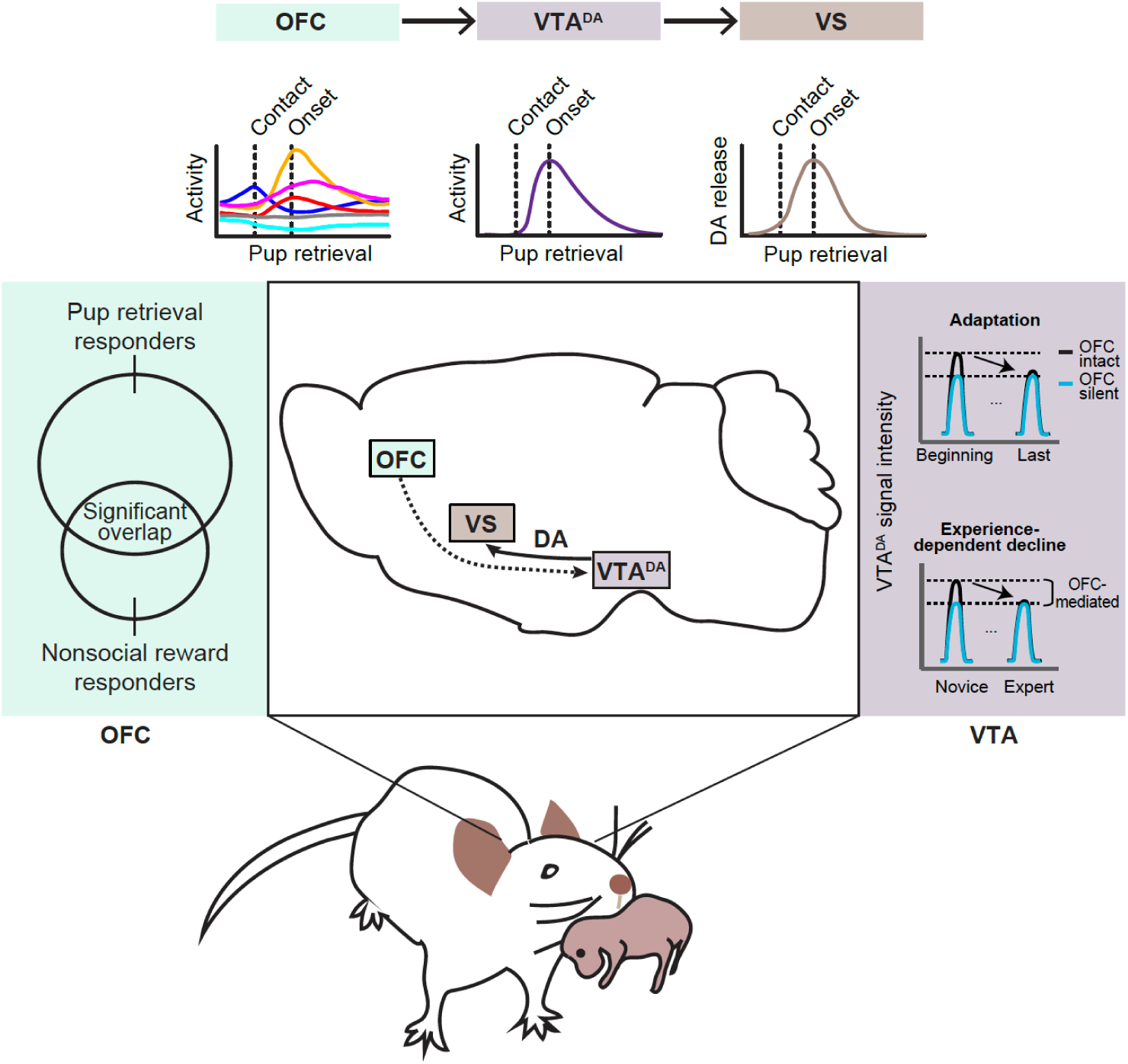
Graphical abstract. The neural representation of pup retrieval in OFC^Rbp4^ neurons significantly overlaps with those responding to nonsocial reward and covers the entire behavioral sequence. VTA^DA^ neurons, which are downstream targets of OFC neurons, show phasic activity peaks at the onset of the pup retrieval, driving a peak in DA release in the VS. Impairment of OFC activity disrupts the dynamic modulation of VTA^DA^ neuron activity at multiple time scales. We propose that OFC activity provides facilitatory signals to VTA^DA^ neurons during the early phase of acquiring alloparental behavior.

**Table S1.**
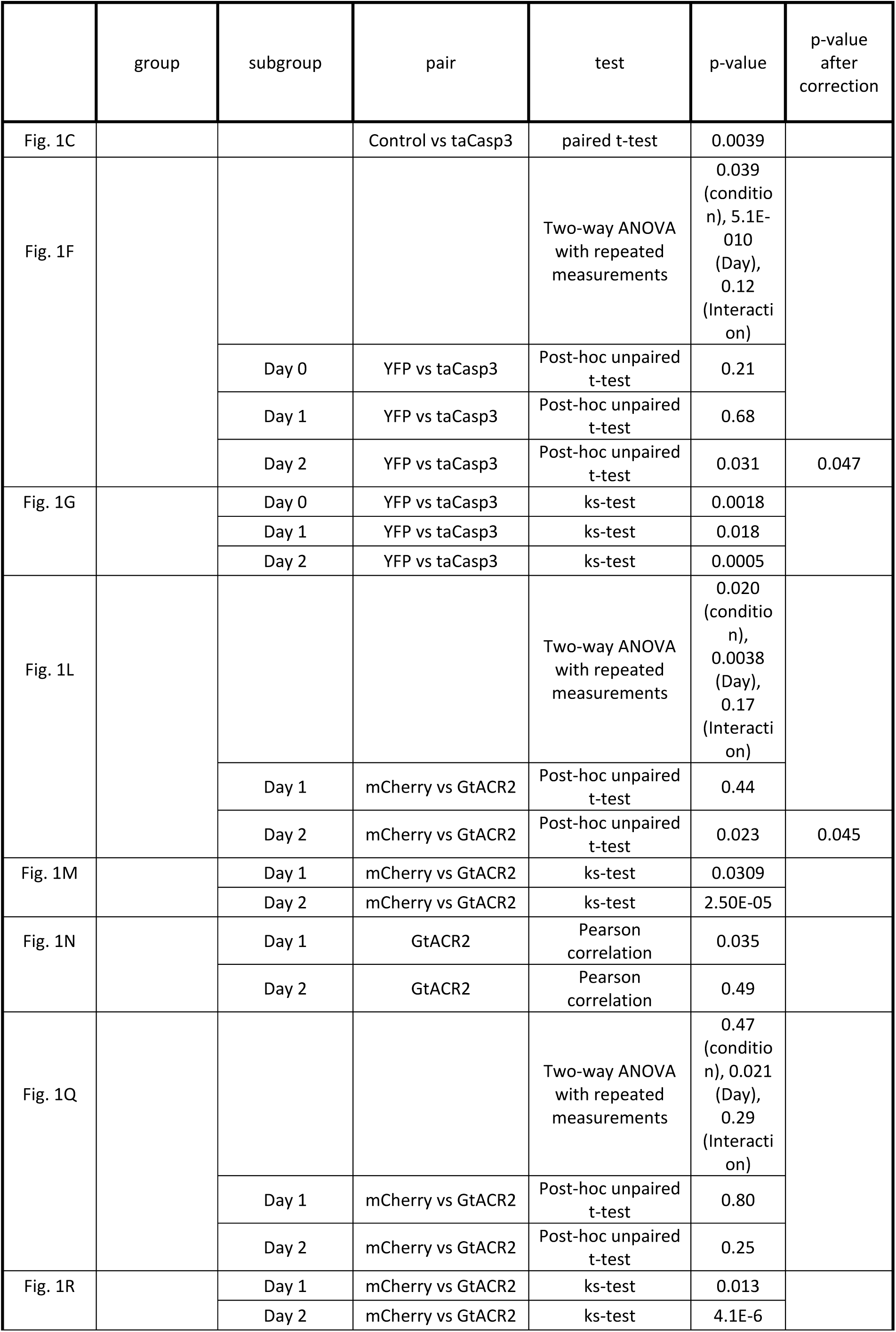
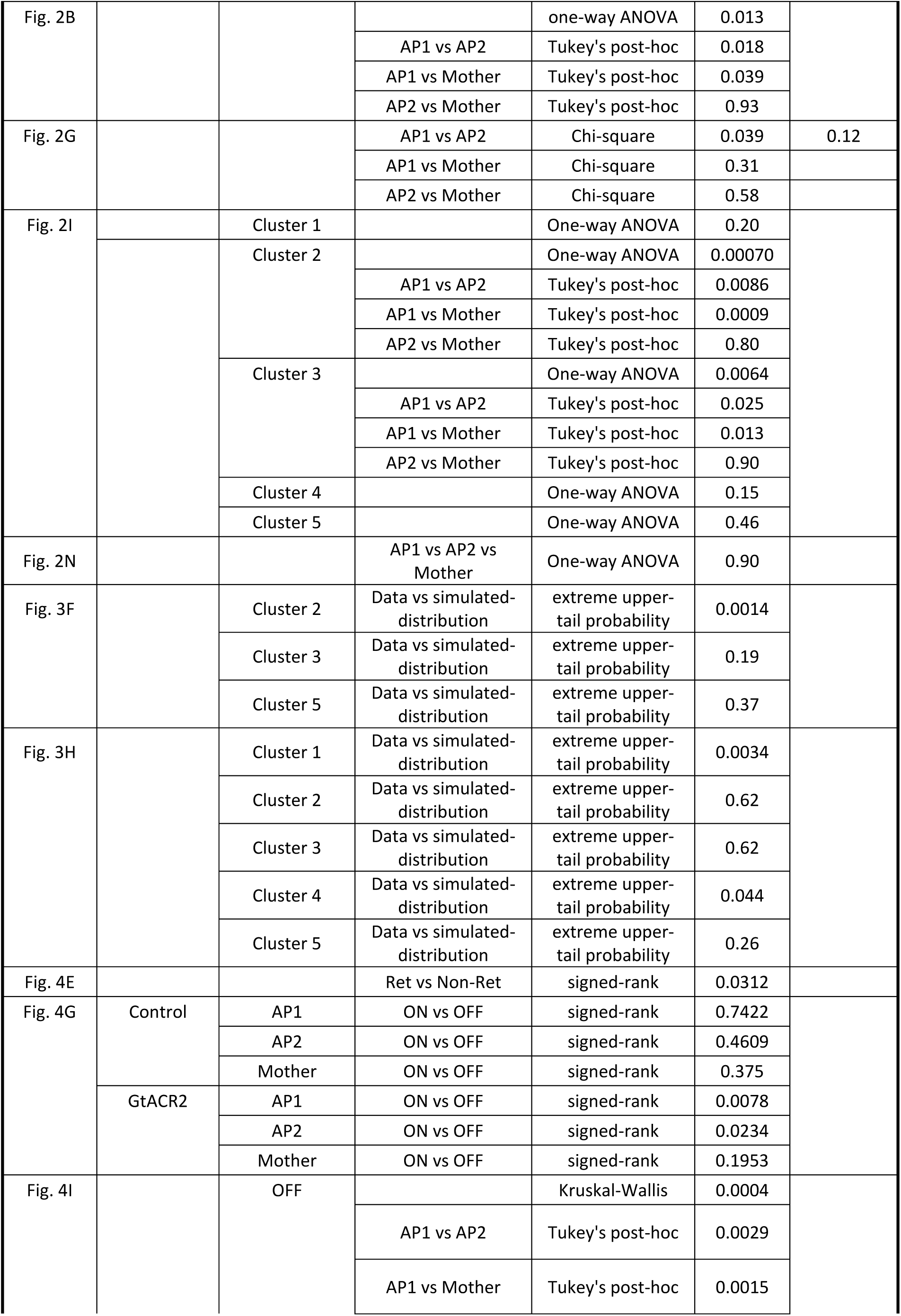
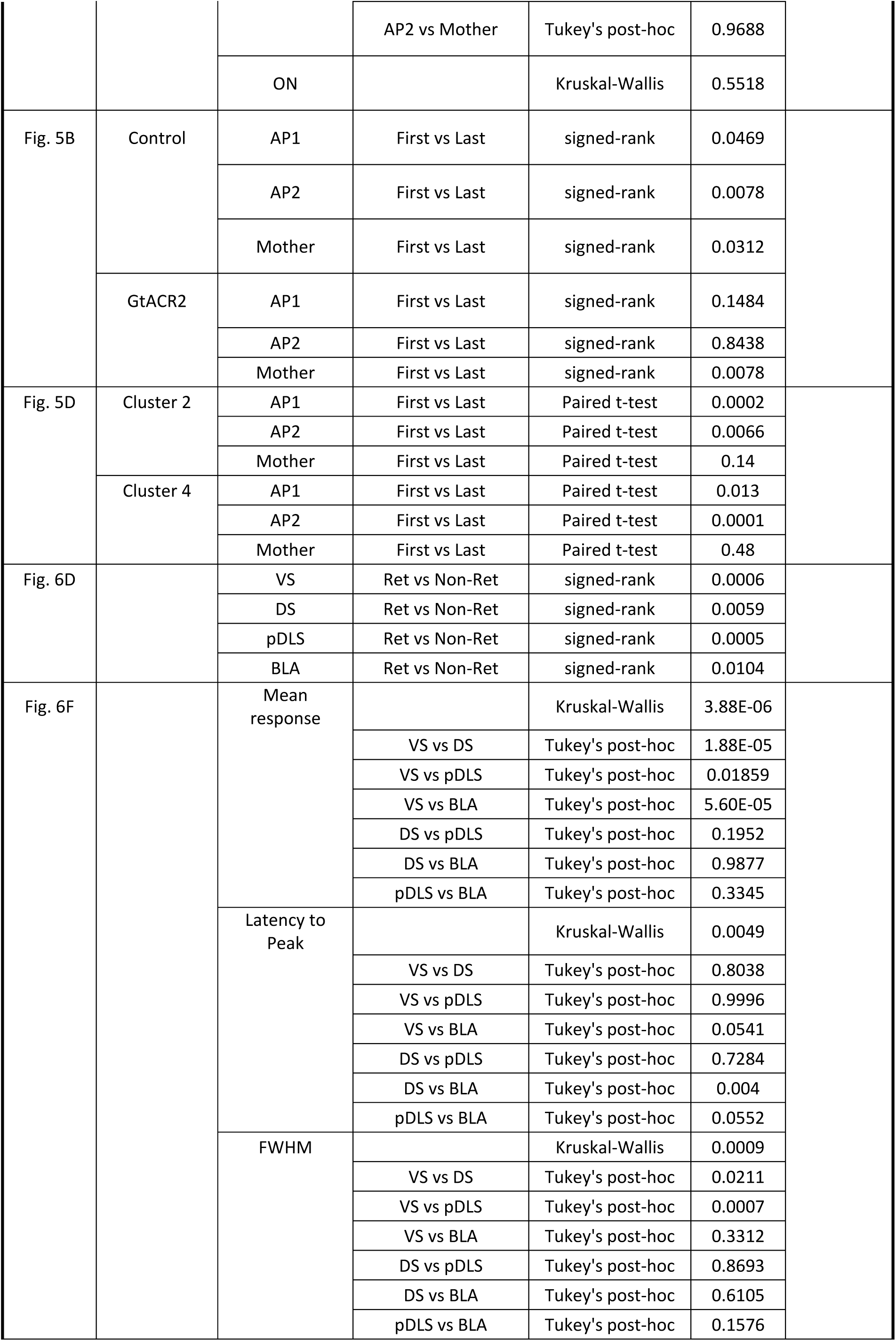
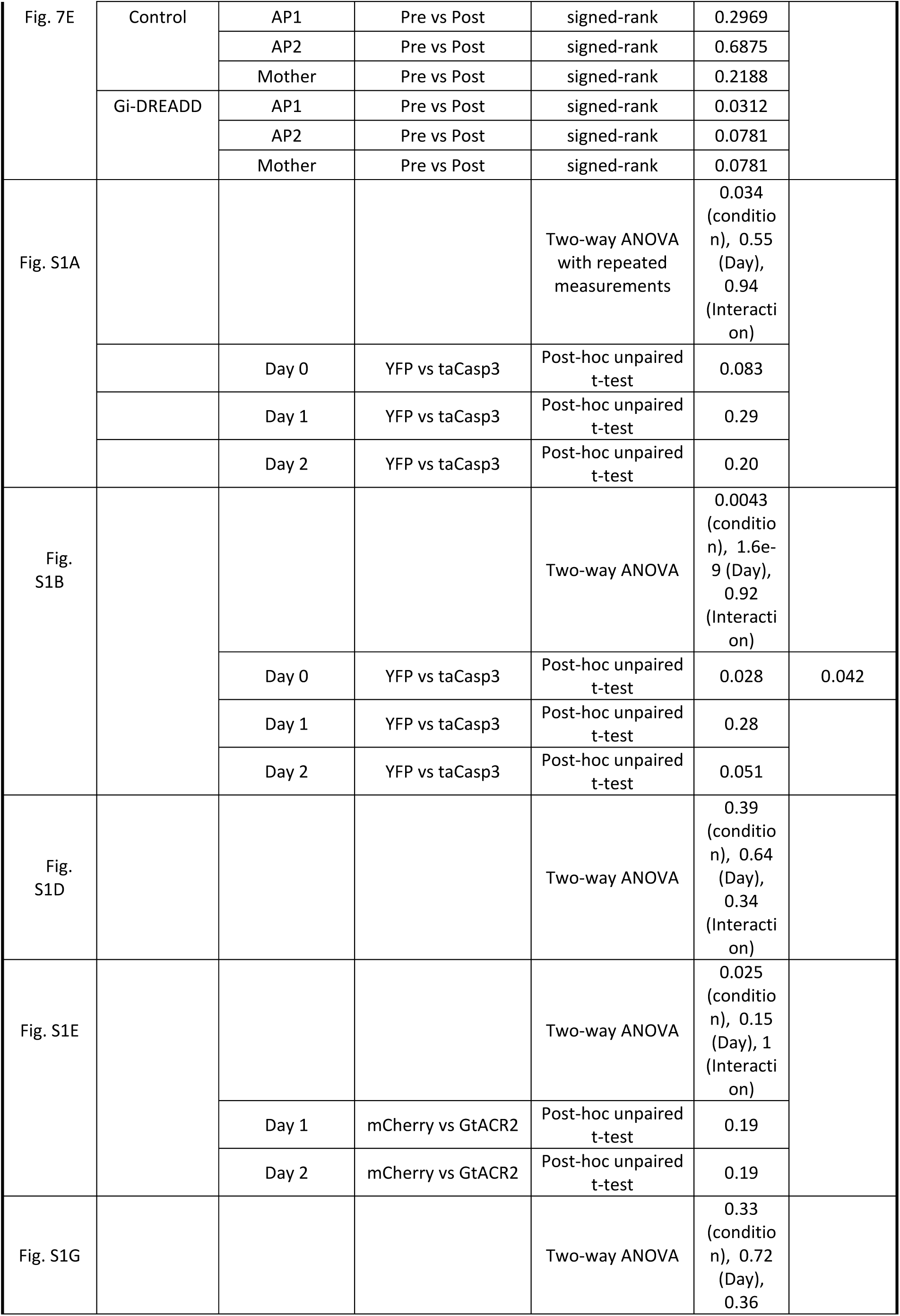
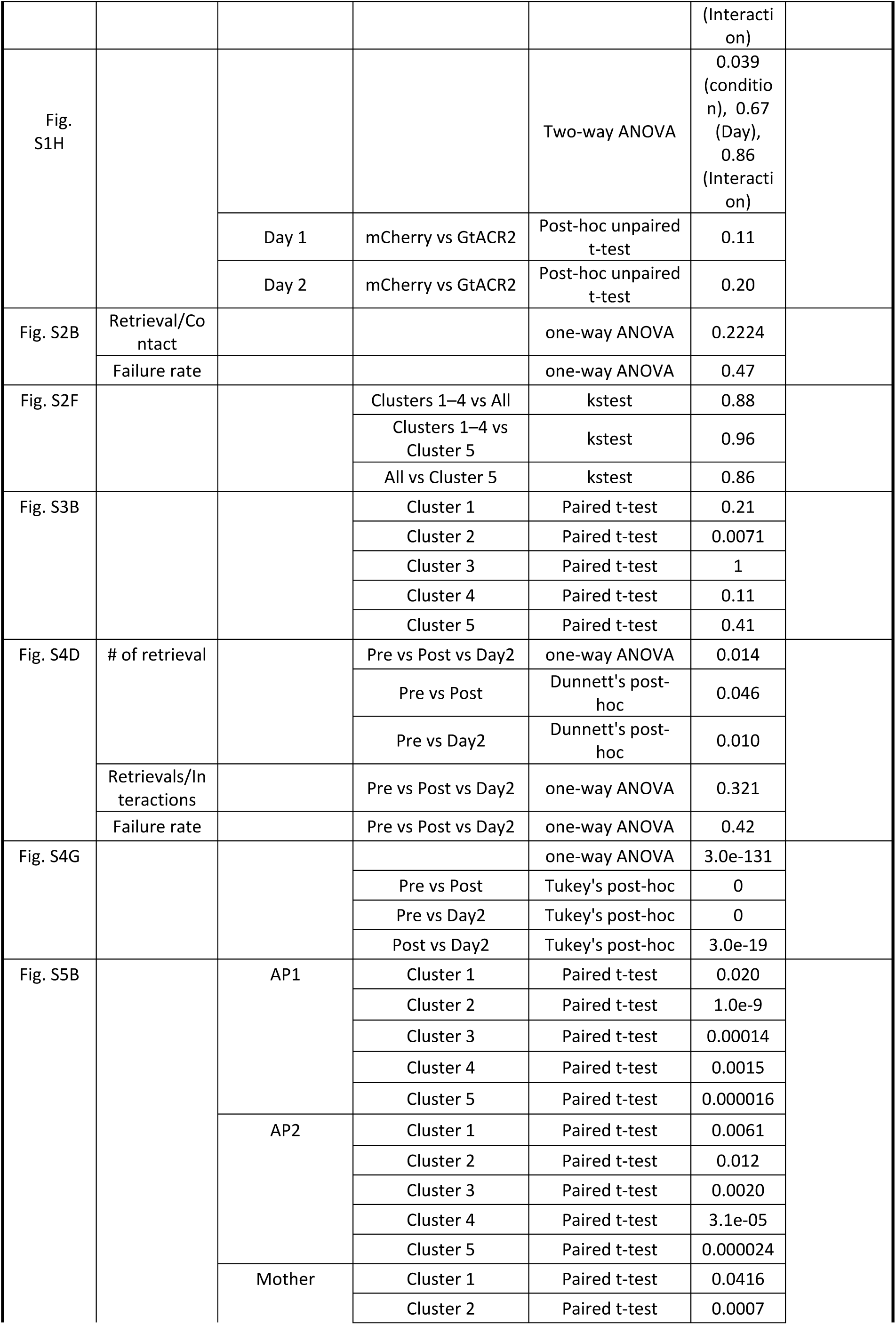
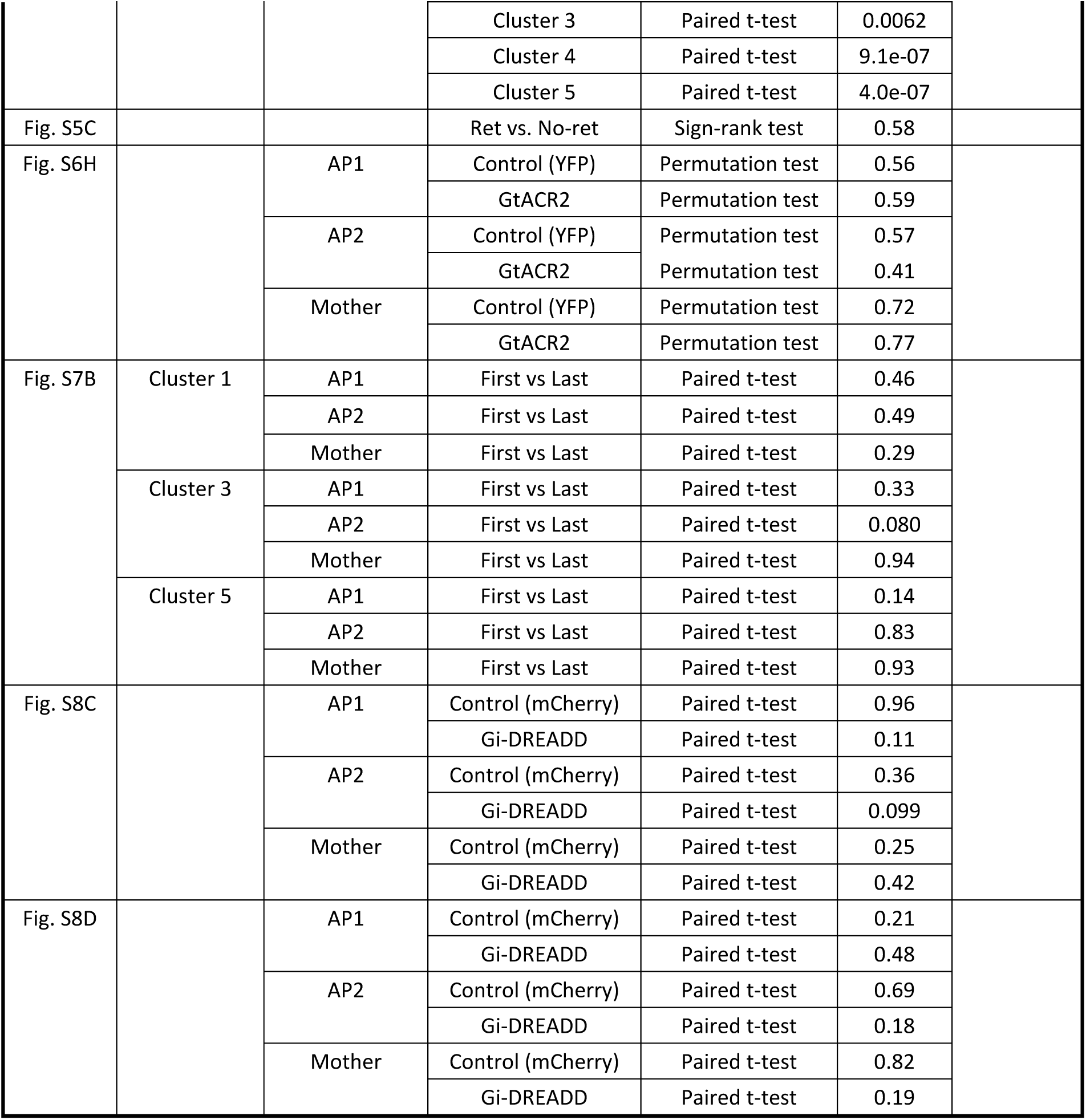
Statistical summary. All tests were two-tailed.

